# Drying halves decomposition rates in river networks by disrupting structure-function linkages

**DOI:** 10.1101/2025.10.07.680619

**Authors:** Rubén del Campo, Thibault Datry, Arnaud Foulquier, Naiara López-Rojo, Loïc Chalmandrier, Zoltán Csabai, David Cunillera-Montcusí, Amaia Angulo, Francisco J. Peñas, Bálint Pernecker, Núria Bonada, José Barquín, Annika Künne, Romain Sarremejane, Maria Soria, Edurne Estévez, Petr Pařil, Heikki Mykrä, Marek Polášek, Barbora Loskotová, Luka Polovic, Gabriel Singer

## Abstract

River drying is intensifying worldwide due to climate change and water abstraction, with major consequences for biodiversity and ecosystem functioning. In river networks, drying not only alters local environmental conditions but also disrupts hydrological connectivity, reshaping the movement of organisms and resources at the network scale. Leaf litter decomposition—a key ecosystem function in freshwater systems—is particularly sensitive to changes in the structure of decomposer communities. We hypothesized that spatiotemporal patterns of drying regulate decomposition by altering the diversity and composition of detritivore macroinvertebrates, bacteria and fungi. We combined data from six European drying river networks (DRNs) spanning a wide latitudinal gradient to assess how local drying intensity and regional hydrological connectivity affect decomposition through changes in these decomposer groups. We found that short drying events (≤ six dry days) reduced decomposition rates by up to 50% by shifting the control of decomposition from a balanced contribution of fungi, bacteria, and detritivores to one dominated by dry-tolerant but less efficient bacteria. These community shifts persisted after flow resumption, leading to sustained reductions in decomposition even under flowing conditions. Regional connectivity alleviated these negative effects of local drying by facilitating the recovery of more efficient aquatic decomposers through dispersal. However, this effect depended on DRN context. In particular, in southern, more arid DRNs, stronger fragmentation hindered the recovery of decomposer communities after flow resumption. Overall, our results provide mechanistic evidence that spatiotemporal patterns of drying can regulate the linkages between community structure and ecosystem functioning in river networks. As drying events become more frequent and prolonged, increasing disruption of these linkages will impact carbon cycling and energy fluxes in freshwater ecosystems under global change.

## Introduction

Rivers rank among the most biodiverse yet most threatened ecosystems globally (Dudgeon, 2019). One of the most pervasive disturbances affecting fluvial ecosystems is drying. Rivers that experience drying events - i.e., cessation and resumption of surface flow - are not limited to arid and semiarid areas, but widespread across all climates (Datry et al. 2023). More than half of the global river network is currently prone to such intermittent or non-perennial flow (Messager et al. 2021). However, the extent of river drying is expected to increase, as many currently perennial river networks may become *drying river networks* (DRNs) under future climate and land-use scenarios (Döll & Schmied 2012, Mimeau et al. 2025). Drying affects the structure - i.e., community composition and diversity - (Gionchetta et al. 2019a, Chalmandrier et al. 2025), and the functioning of river ecosystems (Truchy et al. 2020, López-Rojo et al. 2024). However, we still ignore whether and how drying may disrupt linkages between the structure and functioning across river networks, as well as ensuing ecological impacts.

Biodiversity and ecosystem functioning are intimately linked in river networks (Altermatt 2013, Talluto et al. 2024). Leaf litter decomposition - one of the most essential functions in rivers (Marks 2019) - exemplifies how drying can alter ecosystem functioning by modifying both abiotic conditions and communities. At the local (reach) scale, drying reduces leaf litter decomposition by impairing abiotic processes such as leaching and abrasion, as well as the activity of microbial and macroinvertebrate decomposers. Physiological stress caused by water scarcity constrains decomposer communities (Boulton 1991, del Campo et al. 2021a, Ferreira et al. 2023), often with stronger impacts on detritivorous macroinvertebrates (Leberfinger et al. 2010, Truchy et al. 2020) than on fungi or bacteria (Foulquier et al. 2015, Gionchetta et al. 2019b, Arias-Real et al. 2022). Under intense drying, some detritivores may disappear completely, causing long-lasting legacy effects on decomposition after flow resumption (Datry et al. 2011, Di Sabatino et al. 2022). Moreover, recent evidence suggests that drying can also alter decomposition indirectly through changes in decomposer diversity (Arias-Real et al. 2022). These local-scale effects of drying are well understood, yet it remains unknown whether drying-driven disruptions in hydrological connectivity translate into functional changes at the network scale.

Indeed, at the regional scale, diversity and composition of the principal decomposer groups (bacteria, fungi and detritivores) likely reflect spatiotemporal patterns of drying. Drying events temporarily fragment the network and reduce hydrological connectivity, limiting dispersal and constraining the reassembly of local communities (Gauthier et al. 2021, Jacquet et al. 2022, Sarremejane et al. 2024). When flow resumes, the hydrological reconnection of the network facilitates dispersal, redistributes taxa (Chalmandrier et al. 2025), and can even enhance diversity (Journiac et al. 2025). Thereby, decomposition processes may be re-established through the recovery of decomposer communities in periods of high connectivity following drying (Truchy et al. 2020, Sarremejane et al. 2024). Moreover, changes in hydrological connectivity can also affect decomposition through changes in water physicochemical properties (Bernal et al. 2025), or in the availability of essential nutrients for decomposers (Sarremejane et al. 2024). Overall, restored hydrological connectivity may buffer the impacts of drying on decomposition, while strong fragmentation may amplify them.

As climate and global change create drier, less connected rivers, understanding impacts of drying on ecosystem functioning at both local and network scales becomes crucial. Here, we propose that leaf litter decomposition in DRNs is controlled by shifts in the diversity and composition (i.e., the structure) of decomposers communities at multiple spatial scales, from individual reaches to entire networks. Specifically, our objectives were: (i) to quantify the sensitivity of decomposer communities and decomposition rates to drying, and (ii) to identify how drying alters the constraints of decomposition associated with community structure. We measured leaf litter decomposition rates and sampled microbial and macroinvertebrate communities seasonally at 120 sites across six DRNs along a wide latitudinal gradient in Europe. We hypothesized that: (1) Drying affects decomposition at the network scale through interacting local and regional mechanisms; in particular, we expected high regional hydrological connectivity to buffer negative local impacts through accelerated recovery of decomposer communities (Figure 1). (2) Drying induces long-term shifts in the biotic control of decomposition, from a major contribution of detritivores in perennial reaches to a dominance of microbial decomposers in non-perennial ones, regardless of flowing conditions. Last (3), the strength of drying effects on decomposer communities varies among networks depending on climatic and biogeographical context. By integrating local and regional perspectives, our study advances the mechanistic understanding required to anticipate and mitigate the consequences of climate change and water abstraction for the functional integrity of river networks.

**Figure 1.**
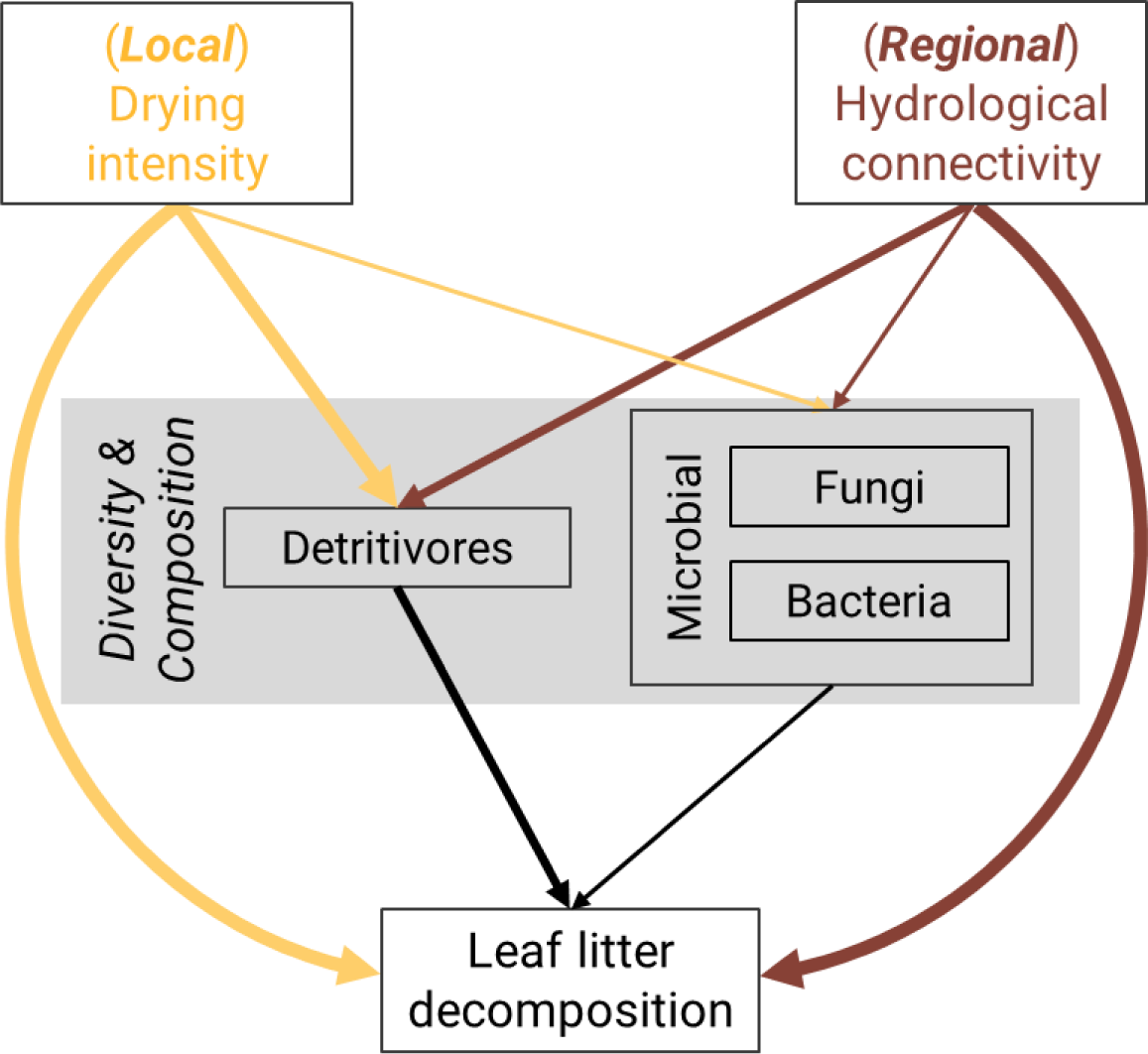
Conceptual path diagram summarising hypothetical local (reach) and regional (network) controls of drying on leaf litter decomposition. Arrows linking drying intensity or connectivity to decomposition via decomposer communities represent indirect effects of drying, mediated by changes in community structure (i.e., composition or diversity) of detritivores, fungi and bacteria. In contrast, the arrows directly linking drying intensity or hydrological connectivity with decomposition represent direct effects of drying not associated with changes in community structure, for instance, abiotic effects or changes in decomposer activity.

## Methods

### Study sites

We sampled 120 reaches across six European DRNs (20 reaches per DRN). Reaches were selected and classified as perennial or non-perennial based on expert knowledge from scientific teams monitoring each DRN, ensuring a balanced representation of both flow regimes. The six DRNs spanned a broad latitudinal gradient covering six ecoregions, from south to north (Figure 2): Genal (GEN) in Spain and Butižnica (BUT) in Croatia in the Mediterranean and Balkan ecoregions, respectively; Albarine (ALB) in France in the Alpine ecoregion; Bükkösdi-víz (BUK) in Hungary in the Pannonian ecoregion; Velička (VEL) in Czech Republic and Lepsämänjoki (LEP) in Finland within the continental ecoregion. All DRNs had similar catchment sizes, ranging from 172 to 354 km^2^. To capture seasonal variability in hydrology, sampling campaigns were conducted in spring (March–June 2021), summer (July–August 2021), and autumn–winter (October 2021–January 2022) (Table S1). During spring sampling, most (84-100 %) reaches across all six DRNs were flowing. In summer, dry reaches peaked in most DRNs (11-43 % dry reaches). In autumn-winter, DRNs were sampled targeting a state after flow resumption, although a considerable proportion of reaches remained dry in some DRNs (0-35 %), reflecting an exceptionally dry year.

**Figure 2.**
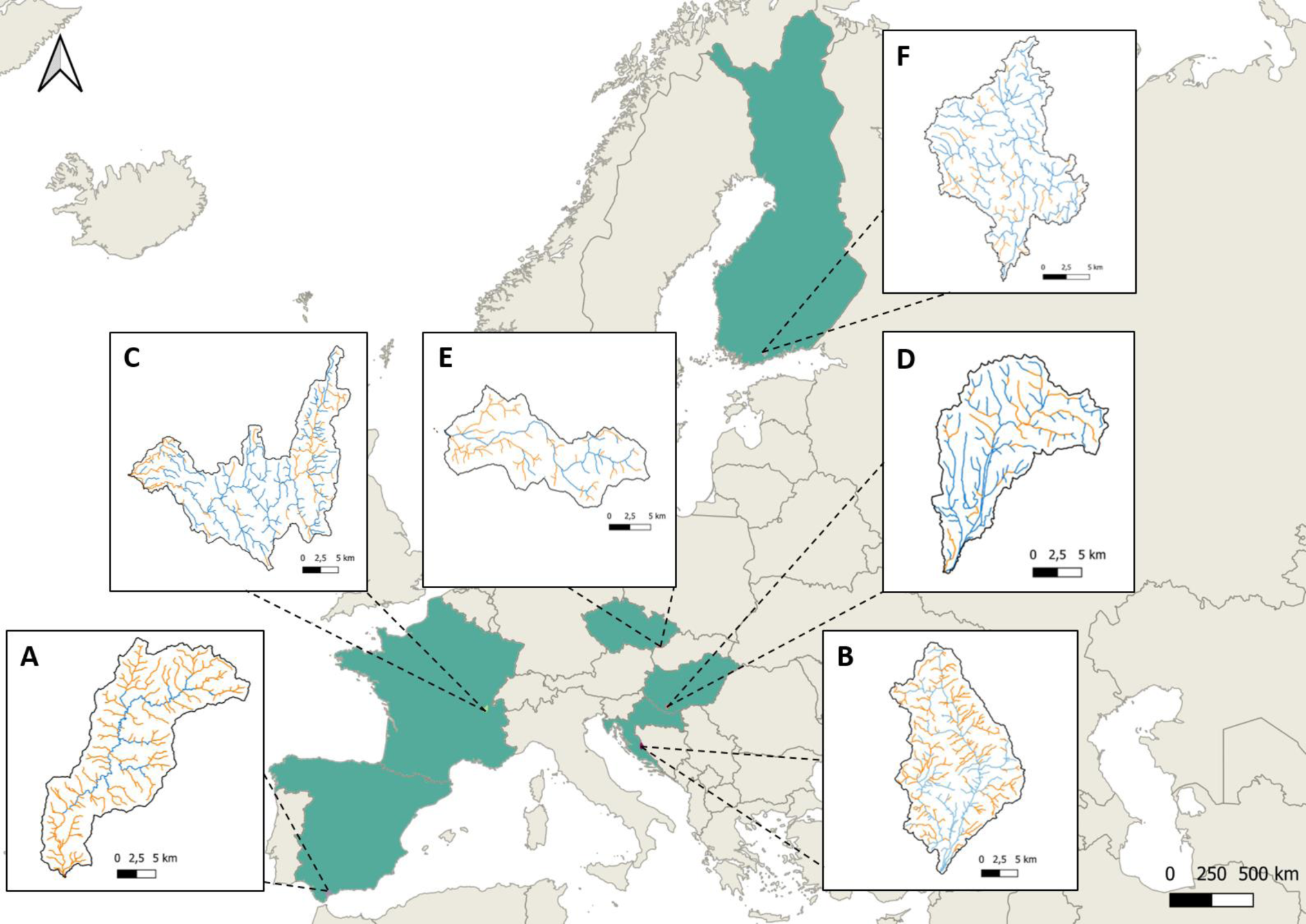
Distribution of the six drying river networks in A) Spain (GEN), B) Croatia (BUT), C) France (ALB), D) Hungary (BUK), E) Czechia (VEL) and F) Finland (LEP). River network reaches are colored based on flow regime (blue = perennial, orange = non-perennial).

### Leaf litter decomposition

For the decomposition experiments, we used three coarse-mesh bags (1 cm opening, 15 x 20 cm size) filled with four grams of air-dried poplar litter, for each sampling reach and campaign. We installed mesh bags either in sections with running water when the reach was flowing, or directly on the dry riverbed when surface flow was absent. At every reach, we deployed a temperature logger (HOBO 8k Pendant, Onset, USA) recording every 30 minutes. Decomposition experiment periods ranged from 8 to 87 days, depending on campaign logistics and aiming at achieving an intermediate decomposition state of leaf litter (20 to 70 % of mass loss). Recollected mesh bags were shipped frozen to the University of Innsbruck, where we processed and measured remaining ash-free dry mass (AFDM) following standard methods (Bärlocher 2020). We estimated temperature-adjusted leaf litter decomposition rates (in units degree-day^-1^) as first-order decay coefficients by regressing the log of remaining AFDM on degree-days as predictor (Bärlocher 2020). No data were available for BUT in summer owing to issues encountered during sample processing.

### Biodiversity sampling

We gathered biodiversity data from microbial and macroinvertebrate communities to characterize community composition and diversity (i.e., the structure) of the principal groups of decomposers, namely detritivorous macroinvertebrates, fungi and bacteria. We sampled biodiversity from January 2021 to February 2022 as close as possible to the decomposition experiments (Table S1). Due to logistical issues, the temporal match between biodiversity samplings and decomposition experiments varied across DRNs and campaigns, ranging from simultaneous samplings (i.e., biodiversity sampling carried out during the decomposition experiment) to a maximum delay of 21 days from the deployment or the collection of the litterbags (Table S1). We discarded reaches for which flow conditions did not match between the biodiversity samplings and decomposition experiments (e.g., a reach that was flowing during the decomposition experiment but dried before sampling biodiversity), yielding 289 observations (237 in flowing reaches, 52 on dry riverbeds).

We sampled macroinvertebrate communities only in flowing sections (disconnected pools were not accounted for) using Surber and/or Hess samplers (500-μm mesh) following a multihabitat approach at each section spanning 50–150m, at least 20 times the average wetted width. Samples were sieved, pre-sorted in the field, and preserved in 96% ethanol, then sorted in the laboratory and identified to the lowest possible taxonomic level (mostly to genus). Abundance data were standardized to individuals/m^2^. Further details can be found in Chalmandrier et al. (2025) and Hárságyi et al. (2025). From the whole macroinvertebrate community, we extracted data corresponding to detritivore taxa. We classified taxa as detritivores when they had an affinity score higher than 1 in the category “dead plant detritus > 1mm” from the food resource trait of the Tachet database, extracted from the Freshwater Information Platform (http://www.freshwaterecology.info) (Schmidt-Kloiber & Hering 2015, Tachet et al., 2010). For taxa not identified to genus or lacking trait information, we assigned an average affinity score based on other genera within the same family. Affinity scores from the fuzzy-coded food trait were converted to percentages. Finally, we calculated the total abundance of detritivores in each sample by multiplying each taxon’s percentage affinity to the “dead plant detritus” category with its abundance (Sarremejane et al. 2024). At dry sites (no surface flow, and thus no macroinvertebrate sampling), detritivore abundance was set to zero. In addition, and only for flowing reaches, we estimated detritivore richness, Pielou’s evenness and changes in community composition using principal coordinate analyses (PCoA). The detritivore PCoA (det.PCoA) was based on Hellinger-transformed abundance and Bray-Curtis dissimilarity. We retained only components explaining more than 2% of the total variance, resulting in seven PCoA components explaining a total variance of 32.23 %. To facilitate the interpretation of PCoA components, we estimated the correlations between the component scores and the relative abundances of each detritivore taxa.

We characterized communities of fungi and bacteria from environmental DNA (eDNA) in sediment samples from dry and flowing reaches using a metabarcoding approach (Taberlet et al., 2012). We collected a sediment composite sample of approximately 0.5 litres from the riverbed to a depth of 10 cm using a gardening shovel or a kicknet, depending on the flowing conditions. This sample was then sieved to remove particles larger than 2 mm. A subsample was stored in sterile 50 ml Falcon tubes and kept frozen at −20 °C pending further metabarcoding analysis. The total extracellular DNA from these samples was extracted using the NucleoSpin® Soil kit (Macherey-Nagel) following Taberlet et al. (2012). The resulting extracts were amplified in triplicate using two sets of tagged markers targeting the 18S gene of Eukaryota, Euka02 (Guardiola et al. 2015) and 16S gene of Bacteria and Archaea, Bact02 (Taberlet et al. 2012). Further details about the DNA extraction and metabarcoding can be found in supplementary material and López-Rojo et al.( 2025). From the 16S and 18S marker tables, we extracted only taxa belonging to the kingdoms Bacteria and Fungi, respectively.

Finally, we removed singletons and calculated MOTUs’ relative abundances using the Hellinger transformation. As for detritivores, we estimated alpha diversity (richness and Pielou’s evenness) and community composition using PCoAs (Bac.PCoA, Fun.PCoA) for bacteria and fungi separately. Bac.PCoA resulted in eight components explaining more than 2 % of variance, explaining a total of 39.23 %. Fun.PCoA resulted in five components, explaining a total of 16.72 %. To better explain Bac.PCoA and Fun.PCoA, we estimated the correlation between the components of each PcoA and the relative abundance of bacteria and fungi MOTUS grouped at class level. To reduce any bias resulting from differences in sequencing depth among samples, we estimated rarefied species richness using the rarefy function from the Vegan package (Oksanen et al. 2020).

### Estimate of drying intensity and hydrological connectivity across DRNs

We characterized two aspects of drying across all DRNs. At the local scale (reach), we estimated the intensity of drying events as the total sum of dry days during the decomposition experiments. We extracted the sequence of dry days during the decomposition experiments from flow intermittence models done for each DRN by Mimeau et al. (2024) using a hybrid modeling approach. At the regional scale, we computed the hydrological connectivity for the 30-days periods preceding each decomposition experiment following the framework established by Cunillera-Montcusí et al. (2023), and using the same networks and link parameters as in Chalmandrier et al. (2025). This approach gives a connectivity index (i.e., spatiotemporal connectivity) that quantifies the relative connectivity between a reach and all its potential network neighbours, ranging between 0 (never connected during the studied period) and 1 (always connected). Chalmandrier et al. (2025) used a temporal window of 30 days before sampling to investigate the response of macroinvertebrate communities to spatiotemporal connectivity. We assumed that a 30-day window would also be appropriate to capture potential influence via microbial communities with shorter life cycles than macroinvertebrates.

### Data analysis

To better understand the control of drying over decomposer communities and decomposition rates, we organized our data analysis in four sequential steps. First, we evaluated differences in decomposition rates between flow regimes and campaigns using mixed models. Second, we examined the capacity of the diversity and composition of bacteria, fungi and detritivores to explain decomposition rates in DRNs through multiple linear regression model selection. Third, we estimated the sensitivity of each decomposer group and of decomposition rates to drying intensity using linear models. And finally, we tested our three hypotheses about the mechanisms controlling decomposition in DRNs (Figure 1) through structural equation modelling (SEM).

The mixed models to test general differences in decomposition rates included flow regime (perennial or non-perennial) and sampling campaigns (spring, summer, winter) as fixed factors, and DRN ID as a random factor. We used the “lme4” R Package to build the models (Bates et al. 2015) and the “emmeans” package (Lenth 2024) to run Tukey contrasts as post-hoc analyses.

Second, we identified and summarized the main diversity and composition features of bacteria, fungi, and detritivores controlling decomposition rates using multiple linear regression models and model selection (MuMIn R package, Bartón 2024). Each group was analysed independently and characterized by its richness (rarefied richness for bacteria and fungi), the evenness of the community (Pielou’s index), and community composition summarised by five to eight PCoA components (see Table 1). The total abundance of detritivores (individuals / m^2^) was also included within the detritivore predictor set. Model selection was based on the Bayesian information criterion (BIC). We retained all models with ΔBIC < 2, and selected the one with the lowest BIC as the most parsimonious model. Finally, for each decomposer group, we used standardized coefficients and 95% confidence intervals from predictors included in the most parsimonious model to represent the magnitude and direction of their effects. The normality and homogeneity of residuals were evaluated using residual plots. We used the most parsimonious model for bacteria, fungi, and detritivores to construct three composite variables, which summarise the influence of the diversity and/or composition (i.e., the structure) of each group on decomposition in our SEMs (see below).

Composite variables (see Grace & Bollen 2008) are multivariate constructs that capture the combined effect of several measured predictors, each contributing complementary information. In practice, composites are estimated as weighted sums of predictors in multiple regression models (e.g., composite = b1*predictor1 + b2*predictor2). While we used the bacterial and fungal composites in all subsequent analyses, we opted to use detritivore abundance instead of the detritivore composite in some cases to include observations from both flowing and dry conditions. Under dry conditions, detritivore abundance can be quantified as zero, while no data on their composition or diversity are available.

Third, we used univariate linear models to estimate the sensitivity to drying intensity (number of dry days) of decomposition rates, detritivore abundance, and the composites for bacteria and fungi. Since we expected a non-linear decline in decomposition rates and detritivore abundance with increasing dry days, we log-transformed both response and predictor variables. A log(x + 1) transformation was applied to the number of dry days and detritivore abundance to account for zero values. Additionally, we used prediction intervals from the linear models to identify drying thresholds, that is, the number of dry days when the response variable dropped below 50% of its maximum value (estimated as the average of the five highest observations during flowing conditions, i.e., zero dry days).

Finally, we used SEM models to test our three specific hypotheses following our conceptual framework (Figure 1). Our first model (SEM1) evaluated the first hypothesis about the interacting effects of local drying intensity and regional hydrological connectivity on decomposition. SEM1 included both direct and indirect links from drying intensity and hydrological connectivity to decomposition rates. Indirect links connected each of the two drying metrics with decomposition rates through detritivore abundance and the composite variables for bacteria and fungi. Thus, indirect links represent drying effects on decomposition associated with changes in the composition and/or diversity of the decomposer community. In contrast, direct links between drying metrics and decomposition rates could be associated with other impacts of drying on decomposition not measured in our study (e.g., abiotic effects, changes in microbial activity, or detritivore activity). In addition, as we expected an influence of drying not only on the abundance of detritivores, but also on their composition and/or diversity, we repeated SEM1 but using the detritivore composite and only flowing reaches (SEM1.1). Combining the results of SEM1 and SEM1.1, we provide twofold evidence about (i) the immediate effects happening along the transition from flowing to dry conditions (SEM1, were both flowing and dry observations are included), and about (ii) long-term effects of drying still persisting after flow resumption (SEM1.1, with only flowing conditions), following our second hypothesis. To provide further evidence on this second hypothesis, we modified SEM1 to run a multigroup SEM with flow regime as a grouping factor (SEM2). Multigroup SEM allows us to evaluate if the relationships among predictors and the response variable vary depending on distinct groups. Thus, we used SEM2 to evaluate whether the control exerted by bacteria, fungi, or detritivores on decomposition rates change between perennial and non-perennial reaches. Since perennial reaches were always flowing (i.e., zero dry days), we constrained the effect of drying intensity only to the non-perennial group. Finally, to address our third hypothesis, we ran a third multigroup SEM (SEM3) with DRN ID as the grouping factor to evaluate potential differences in the mechanisms of drying control that could emerge across the studied DRNs. SEM models were built using the package Lavaan (Roseel, 2012). To improve linearity, we log-transformed both decomposition rates and dry days(log + 1 for dry days), and square root-transformed hydrological connectivity. Because strong correlations among predictors can impair SEM performance, we accounted for expected covariance among decomposer groups, by including correlation errors between composite variables and/or detritivore abundance in all SEMs as necessary. Covariance could come either from interactions among groups, or due to common reactions to drying. Model fit and performance were evaluated using the modification index test, Akaike Information Criterion (AIC), the root mean square error of approximation (RMSEA), comparative fit index (CFI), and standardized root mean squared residuals (SRMR). The fitting power of the model (R-square) was measured based on the amount of explained variation of decomposition rates. All analyses and plots were performed with R software, version 4.3.2 (R Core Team 2023).

**Table 1.** Summary of all predictors (1) included in model selection to characterize the composition and diversity of each decomposer group, and (2) selected in the most parsimonious model for each group. PCoA components are described based on the sign of their correlation with associated taxa or MOTUS.

| 1 - All predictors used for model selection to create composite variables |  |  |
| --- | --- | --- |
| Group | Predictor | Definition |
| All | Richness | Number of taxa (richness), rarefied in case of fungi and bacteria |
| All | Community evenness | Pielou's index |
| All | Community composition | PCoA components based on Hellinger-transformed abundance. Only components explaining more than 2 % of total variance. |
| Detritivores | Abundance | Total density of detritivores (individuals / m<sup>2</sup>) at sampling reach. Abundance equal to 0 in dry reaches. |
| 2 - Predictors selected in best models |  |  |
| Group | Predictor | Correlations of taxa / MOTUS with PCoA components |
| Bacteria | Bac.PCoA1 | corr -: Actinobacteria, Thermoleophilia.<br>corr +: Beta- and deltaproteobacteria, Bacteroidia, Verrucomicrobiae |
|  | Bac.PCoA2 | corr -: Gammaproteobacteria, Bacteroidetes<br>corr +: Actinobacteria, Acidobacteria |
|  | Bac.PCoA3 | corr -: Acidobacteria, Alphaproteobacteria, Deinococci<br>corr +: Actinobacteria, Longimicrobia, Thermoplasmata, Cytophagia |
|  | Bac.PCoA6 | corr -: Acidimicrobia, Actinobacteria, Alphaproteobacteria, Chloroflexi<br>corr +: Nitrospira |
|  | Bac.PCoA8 | corr -: Actinobacteria, Gemmatimonadetes<br>corr +: Acidobacteria, Terrimicrobia |
| Fungi | Richness |  |
|  | Evenness |  |
|  | Fun.PCoA1 | corr -: Chytridiomycetes<br>corr +: Pezizomycetes, Agaricomycetes, Ascomycota (undefined class) |
|  | Fun.PCoA2 | corr -: Leotiomycetes, Sordariomycetes<br>corr +: Agaricomycetes, Pezizomycetes |
| Detritivores | Abundance |  |
|  | Det.PCoA5 | corr -: Leuctra, Nemoura, Echinogammarus, Nigrobaetis, Sericostoma, Rhyacophila<br>corr +: Tabanidae, Caenis |
|  | Det.PCoA7 | corr -: Limoniidae, Ecdyonurus, Leuctra, Electrogena, Sericostoma, Habroleptoides<br>corr +: Nemoura, Baetis |

## Results

### Summary of drying and decomposition patterns across DRNs

Local and regional drying patterns varied greatly across the six DRNs (Table S2, Figure S1). LEP and ALB networks experienced milder drying periods, with the lowest values of drying intensity (median of 0 and 5 dry days, respectively; Figure S1a) and the highest values of hydrological connectivity (median of 0.98 and 0.45, respectively; Figure S1b). GEN and BUT - the southernmost DRNs - were characterized by the lowest values of hydrological connectivity (median of 0.07 and 0.13, respectively), and BUT registered also the longest dry periods (median of 43 dry days). BUK and VEL showed high drying intensity (median of 11 and 15 dry days, respectively) and intermediate connectivity levels (median of 0.23 and 0.24, respectively).

Leaf litter decomposition rates varied between 10 to 96 % mass loss, corresponding to decomposition rates of 3.5 x 10^-4^ and 7.3 x 10^-3^ dd^-1^. Perennial reaches had significantly higher decomposition rates than non-perennial ones (F_1,278_: 80.55, p < 0.001; Figure 3a). Differences in decomposition between flow regimes persisted across all campaigns (Tukey t-ratio ranging from -2.8 to -7.7, p < 0.01; Figure 3b). However, a significant interaction between flow regime and campaign (F_2,278_: 7.50, p < 0.001) indicated that differences across campaigns were only significant for non-perennial reaches (Tukey t-ratio ranging from -2.3 to 5.6, p < 0.05), where decomposition rates dropped sharply during the summer due to drying (Figure 3).

**Figure 3.**
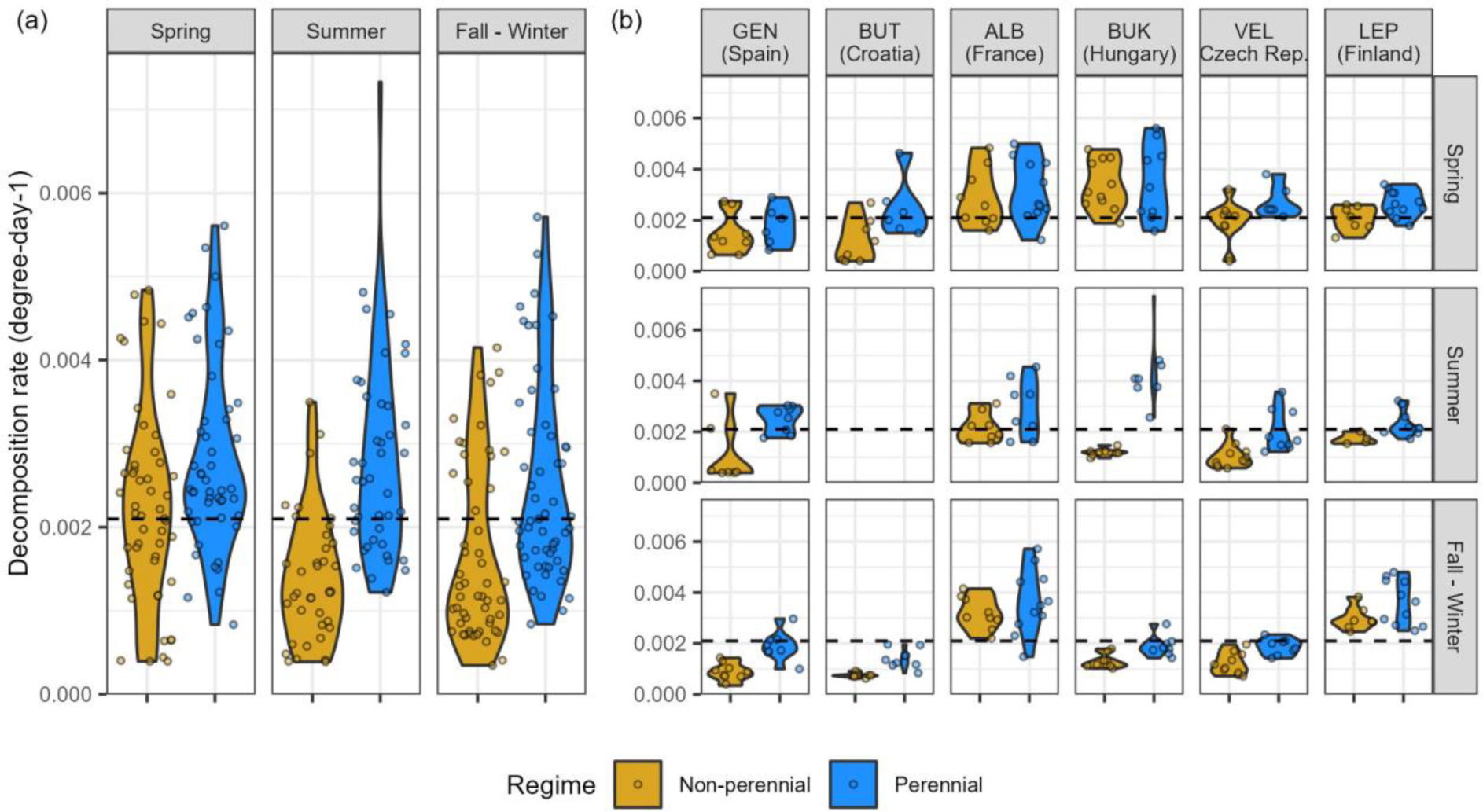
Violin plots representing the variability of leaf litter decomposition rates among (a) flow regimes and sampling campaigns, and (b) depending on the DRN. The horizontal dashed line represents the median value of decomposition rates for the whole dataset (k = 0.0021 degree-day^-1^). No data were available for BUT in summer owing to issues encountered during sample processing.

### Model selection and creation of composite variables for each decomposer group

Model selection identified a single candidate model (ΔBIC < 2) explaining the relationship between bacterial community composition and diversity with decomposition rates (Table S3). The most parsimonious model (F₅,₂₆₅ = 61.3, p < 0.001, R² = 0.53) included five PCoA components summarising bacterial community composition (Bac.PCoA1, Bac.PCoA2, Bac.PCoA3, Bac.PCoA6, and Bac.PCoA8; Table S3, Figure S2). Negative bacterial composite values corresponded to low decomposition rates occurring in dry reaches dominated by the classes Actinobacteria, Thermoleophilia, and Longimicrobia (i.e., negative scores of Bac.PCoA1 and positive scores of Bac.PCoA3, Figure S3). Positive composite values corresponded to higher decomposition rates in flowing reaches, where Nitrospira, Beta- and Deltaproteobacteria were more prevalent (i.e., positive scores of Bac.PCoA1, Bac.PCoA6, and Bac.PCoA8; Figure S3).

For fungi, two candidate models were found (ΔBIC < 2, Table S3). The most parsimonious model (F_4,258_ = 31.5, p < 0.001, R^2^ = 0.32) included the Fun.PCoA1 and Fun.PCoA2 components, the richness and the evenness of the community (Table S3, Figure S2).

Negative values of the fungal composite were associated with low decomposition rates in dry reaches dominated by the classes Leotiomycetes, Sordariomycetes, Pezizomycetes and some MOTUS of Ascomycota with undefined class (positive scores of Fun.PCoA1, negative scores of Fun.PCoA2; Figure S4). Positive composite values were associated with high decomposition rates in flowing reaches with higher richness, higher evenness and high relative abundance of Chytridiomycetes (i.e., positive scores of Fun.PCoA2; Figure S4).

For detritivore communities in flowing conditions, we identified five candidate models (Table S3). The most parsimonious model (F_3,222_ = 16, p < 0.001, R^2^ = 0.17) included the detritivore abundance, Det.PCoA5 and Det.PCoA7 components (Table S3, Figure S2). Positive values of the detritivore composite were associated with high decomposition rates, mainly related to high abundances of detritivores, particularly *Gammarus* and *Nemoura* (i.e., positive scores det.pcoa7; Figure S5). Negative detritivore composite values were associated with lower decomposition rates found in reaches with lower detritivore abundance, and dominated by Diptera taxa (i.e., positive scores det.pcoa5, negative scores PCoA7; Figure S5).

### Sensitivity of decomposition rates and decomposer communities to drying intensity

Decomposition rates, detritivore abundance, and the composite variables summarising community composition for bacteria and fungi, all decreased exponentially with increasing drying intensity (Figure 4). Linear models predicted a 50% drop in decomposition rates after 6.1 dry days (range 4.6 - 8.5 dry days, F_1,287_ = 166, R^2^= 0.36; Figure 4a). From all decomposer groups, detritivores showed the strongest sensitivity to drying, reaching a 50 % drop in abundance just after 1.8 dry days (range 1.4 - 2.2 dry days, F_1,277_ = 313, R^2^= 0.53; Figure 4b). Fungi and bacteria were less sensitive to drying, reaching the 50 % drop in composite values after 13 and 18.3 dry days, respectively (Fungi: range 8.7 - 21.9 dry days, F_1,261_ = 111, R^2^= 0.30; Bacteria: range 13.1 - 27.5 dry days, F_1,261_ = 204, R^2^= 0.44; Figure 4c, Figure 4d).

**Figure 4.**
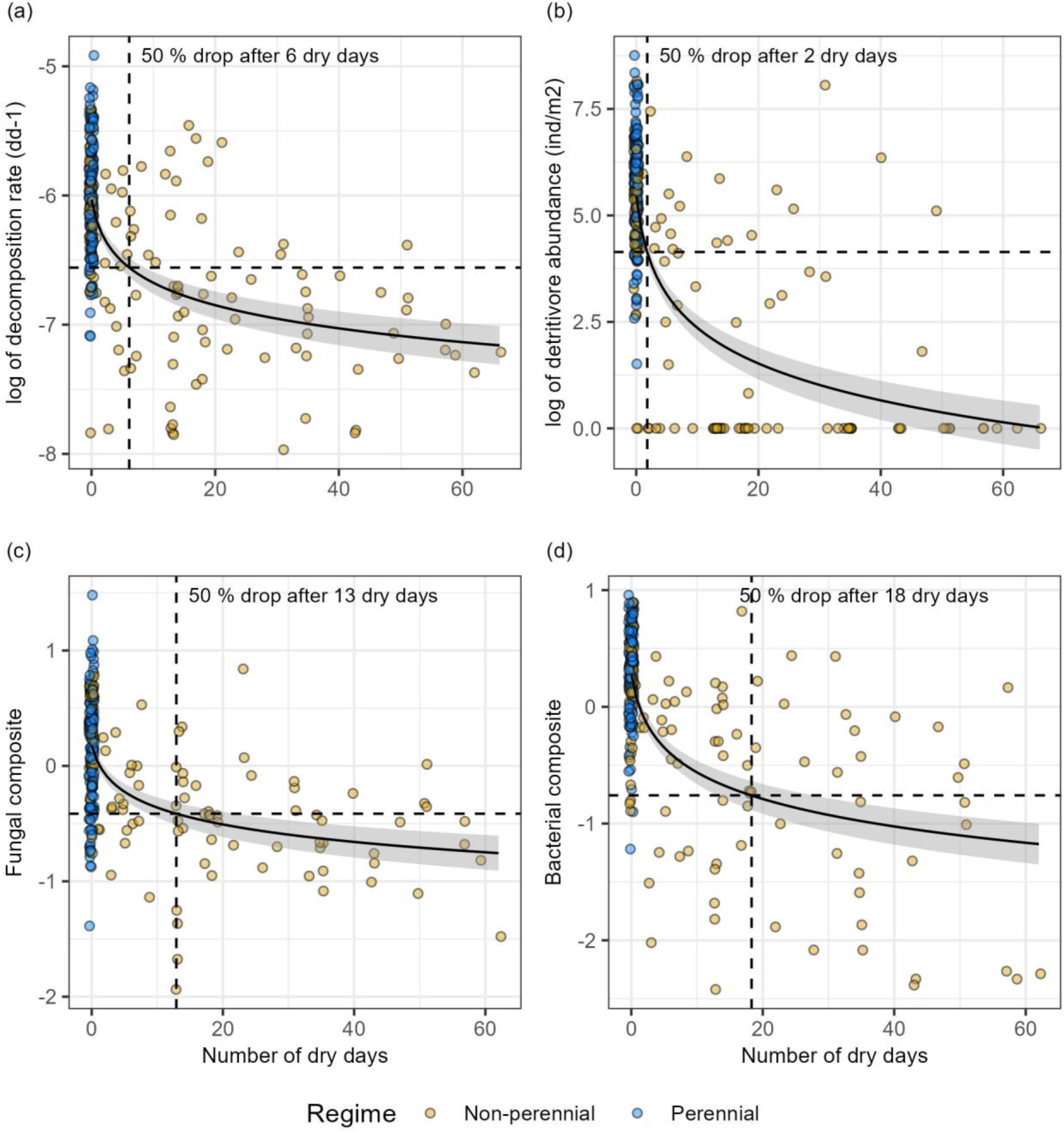
Linear models used to estimate the sensitivity to drying intensity of (a) litter decomposition rates, (b) detritivore abundance, (c) bacterial, and (d) fungal composites. The point where vertical and horizontal dotted lines meet indicates the number of dry days at which the study response drops below 50 % from its maximum value (average of the five highest values at zero dry days). Grey areas represent 95% prediction intervals.

### SEM models

SEM1 showed a satisfactory fit, indicating no discrepancy between the proposed model and the data (chi-square = 1.59, 1 df, p = 0.207; Figure 5, Table S4). The model explained litter decomposition rates with an R^2^ = 0.62, detritivore abundance with an R^2^ of 0.55, the bacterial composite with an R^2^ = 0.45, and fungal composite with an R^2^ = 0.35. Following our first hypothesis, SEM1 revealed a significant indirect and negative effect of drying intensity on decomposition rates at the local scale, but a positive influence of hydrological connectivity at the regional scale (Figure 5). The direct link between drying intensity and decomposition rates was not significant. Instead, increased drying intensity led to a marked decline in detritivore abundance and pronounced changes in fungal and, especially, bacterial composites. The negative effect of drying on both bacterial and fungal composites suggested a replacement of taxa typical of flowing conditions by those more commonly associated with dry conditions. Hydrological connectivity exerted a significant positive influence on decomposition rates through both direct and indirect links. Community-mediated effects (indirect links) were mainly mediated through a positive effect on both bacterial and fungal composites, with a comparatively stronger effect on fungi. Connectivity also exerted a positive but non-significant effect on detritivore abundance. Finally, SEM1 revealed a much stronger control of bacteria on decomposition rates in DRNs, than either fungi or detritivore abundance. The strong covariance between decomposer groups was accounted for by correlation errors between the bacterial and fungal composite, and between the bacterial composite and detritivore abundance.

**Figure 5.**
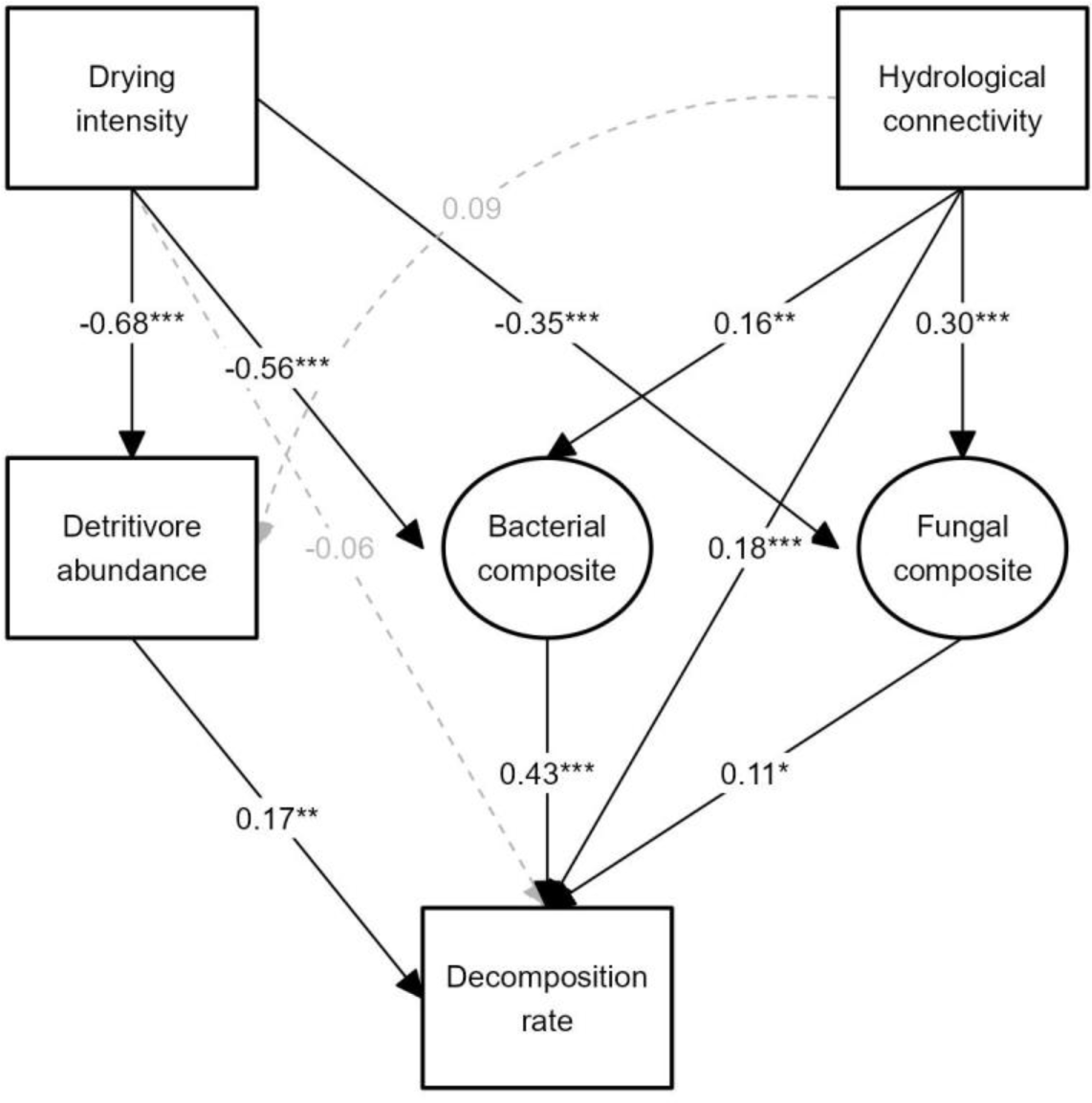
Structural equation model (SEM1) describing the local (drying intensity) and regional (hydrological connectivity) mechanisms controlling leaf litter decomposition rates in DRNs. Boxes represent measured parameters, while the circles represent composite variables created to summarise the effect of bacterial and fungal community structure (composition and/or diversity) on litter decomposition. Black solid lines and associated numbers represent significant paths and standardised path coefficients. Grey dashed lines and numbers represent non-significant paths.

The model SEM1.1 also resulted in a satisfactory fit to the data (total chi-square = 0.048, df = 1, p = 0.826; Table S3) indicating that the effects of drying on decomposition rates persisted during flowing conditions, following our second hypothesis (Figure S6). In SEM1.1, drying intensity decreased decomposition rates during flowing conditions through a negative effect on the detritivore composite. This negative effect was mediated by the replacement of detritivores with high abundances, such as *Gammarus* or *Nemoura* by Diptera taxa, such as Tipulidae, Limoniidae, or Ceratopogonidae (Figure S5). In contrast to SEM1, hydrological connectivity did exert a significant, positive effect on the detritivore composite in SEM1.1 (Figure S6), indicating a stronger link between regional connectivity and community composition than with the total abundance of detritivores. In agreement with SEM1, bacteria emerged as the most important group controlling decomposition rates, showing path coefficients twice as high as of the detritivore composite, and four times greater than of fungi. To test further our second hypothesis, we carried out a multigroup analysis with flow regime as grouping variable (SEM2). SEM2 showed a good fit to the data (total chi-square = 4.494, df = 2, p = 0.106; Table S3) and clearly distinguished biotic controls of decomposition in perennial and non-perennial reaches (Figure 6). In perennial reaches, decomposition rates were equally controlled by the three decomposer groups (i.e., similar path coefficients for detritivore abundance, bacterial and fungal composites). In contrast, in non-perennial reaches, only the bacterial composite exerted a significant effect on decomposition rates, with more than five times greater influence on decomposition than fungi or detritivore abundance. Regional connectivity had a significant, positive effect on all decomposer groups in non-perennial reaches. In contrast, connectivity only affected fungi in perennial reaches.

**Figure 6.**
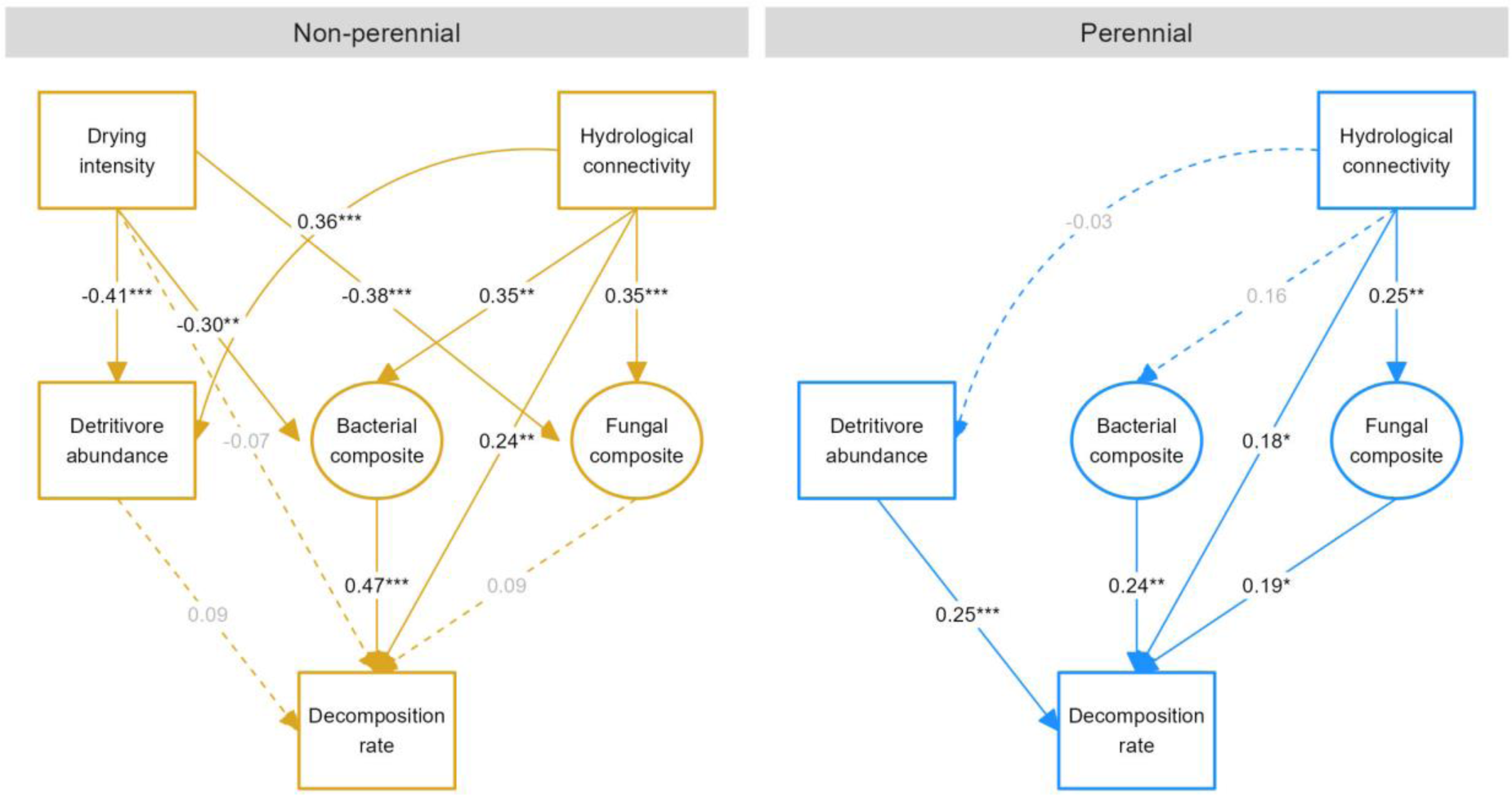
Multigroup structural equation model (SEM2) describing differential biotic controls of decomposition rates in DRNs depending on flow regime. Boxes represent measured parameters, while the circles represent composite variables created to summarise the effect of bacterial and fungal community structure on litter decomposition. Solid lines and associated numbers represent significant paths and standardised path coefficients. Dashed lines with grey numbers represent non-significant paths.

Finally, the multigroup model SEM3 using DRN as a grouping factor showed also a satisfactory fit to the data (total chi-square = 12.006, df = 6, p = 0.062; Table S3). In agreement with our general model (SEM1), all DRNs showed a negative effect of drying on decomposition rates through the alteration of, at least, one decomposer group. Yet, the in-depth analysis by DRN showed certain differences in the biotic mechanisms controlling decomposition depending on the DRN (Figure S7). We found that the effect of hydrological connectivity on decomposition, as well as the contribution of the various decomposer groups, changed with DRN following a south to north gradient. Hydrological connectivity had no significant effects on decomposition rates in southernmost DRNs (GEN, BUT and ALB).

There, decomposition rates were mainly controlled by bacterial communities, characterized by negative scores of the composite (Figure S8b). In central European and northernmost DRNs (BUK, VEL and LEP), hydrological connectivity had a stronger influence on decomposer groups, and/or directly on decomposition rates. Decomposition rates were mainly controlled by detritivores in BUK and VEL, where they reached maximum abundance (Figure S8a). In LEP, decomposition rates were mainly associated with bacterial and fungal communities, characterized by positive composite values (Figure S8).

## Discussion

Here we show for the first time how regional connectivity and local drying events interact to control decomposition patterns across six river networks spanning a broad latitudinal gradient in Europe. We show that drying can control decomposition rates at the river network scale by altering the diversity and composition of bacteria, fungi and detritivores. In particular, local drying events can halve decomposition rates by replacing efficient but drying-sensitive decomposer communities by dry-tolerant bacteria. However, regional connectivity can alleviate such negative impacts of drying by promoting the recovery of decomposer communities and enhancing the exchange of essential resources (e.g., nutrients) across river networks. Still, the contribution of the various decomposer groups, as well as their response to hydrological connectivity, can change depending on climate and/or biogeography, with more arid networks in southern Europe showing stronger impacts of network fragmentation, hindering the recovery of fungi and detritivores.

### Local drying events halve decomposition rates by disrupting biodiversity-functioning linkages across river networks

Previous studies show that drying reduces decomposition rates either by limiting physical breakdown by water and sediment flow (Ferreira et al. 2023) or due to physiological constraints on microbial activity (Foulquier et al. 2015, Gionchetta et al. 2019b, Arias-Real et al. 2022) and detritivores (Leberfinger et al. 2010, Truchy et al. 2020). Our SEM approach allowed us to differentiate changes in decomposition caused by shifts in diversity and community composition (drying indirect links) from other effects associated with unmeasured factors such as decomposer activity (drying direct link). The direct link between drying intensity and decomposition rates was not significant in most SEMs, thus suggesting a higher importance of structural changes rather than in the activity of decomposers. Furthermore, combining SEMs and threshold analyses, we could not only identify the mechanisms by which drying alters the diversity and composition of decomposer communities, but also quantify their sensitivity. We found that just six dry days were enough to halve decomposition rates (Figure 4a) due to fast structural changes in decomposer communities. According to our results, decomposition rates decrease immediately with the onset of drying due to the fast reduction of detritivore abundance (50 % drop after 2 days of drying). Afterwards, drying induces changes first in the community composition of fungi (50 % change after 13 dry days) and finally bacteria (50 % change after 18 days). These results are in line with previous studies that found that short drying events (e.g., less than 10 days) can disrupt microbial decomposition (Arroita et al. 2018, Truchy et al. 2020, Simoes et al. 2021). Taken together, these findings suggest that within less than a month of drying, the structure-mediated control of decomposition shifts from a balanced influence of the structure of detritivore, bacterial, and fungal communities on litter decomposition (as indicated by comparable SEM path coefficients; Figure 6), to a situation in which decomposition is predominantly driven by the variation in potentially dry-tolerant bacterial communities. Interestingly, our results also show that this re-structuring of the decomposer community not only affects dry reaches, but persisted into the flowing phase of non-perennial rivers (SEM1.1, Figure S6), explaining the overall lower decomposition rates in non-perennial than in perennial reaches (Figure 3).

Based on six river networks, our results indicate that the long-lasting impact of drying on functioning (i.e., the drying legacy effect) previously found in individual reaches (Datry et al. 2011, Di Sabatino et al. 2022, Ferreira et al. 2023), also operates across entire river networks. Such long-term impact at the network scale can be associated with the limited capacity of both microbial (Pohlon et al. 2013, Truchy et al. 2020) and macroinvertebrate communities (Sarremejane et al. 2021, Di Sabatino et al. 2022) to fully recover after a strong disturbance caused by drying. Overall, our findings demonstrate that drying not only alters local decomposer communities or decomposition rates at reach scale, but also disrupts structure-functioning relationships at the river network scale.

### Drying-induced changes in decomposer communities bring a decrease in decomposition rates

The impact of local drying events on decomposition rates have traditionally been associated with the constraint of decomposer activity (Boulton 1991, del Campo et al. 2021a, Ferreira et al. 2023). Here, we propose a supplementary mechanism to explain the decrease of decomposition rates based on the replacement of effective, aquatic decomposers by taxa with higher capacity to handle drying stress, but lower capacity to decompose leaf litter. In a recent study, Arias-Real et al. (2023) showed the existence of a strong functional trade-off between the decomposition efficiency of fungi and their capacity to survive drying. Our results suggest that this trade-off can be extended to the whole decomposer community, from fungi to detritivores and bacteria. For instance, some key detritivore taxa often characterized by very high abundances (e.g. *Gammarus*) also have minimal capacity to deal with flow intermittence (Arias-Real et al. 2021, Pařil et al. 2019); thus, they are among the first taxa to disappear at the onset of drying (Datry et al. 2011, Di Sabatino et al. 2022). We observed that drying intensity promoted the replacement of these efficient taxa by generalists and r-strategists relying on increased resilience, such as Diptera (Figure S5, Figure S6).

Similarly, in communities of fungi and bacteria, we observed that drying events led to a replacement of more aquatic taxa (e.g., Beta- and Deltaproteobacteria or Chytridiomycetes for fungi) by soil-inhabiting assemblages dominated by drying specialists (e.g., Actinobacteria, Thermoleophilia, for bacteria or fungi like Dothideomycetes, Pezizomycetes, Agaricomycetes or Leotiomycetes) as previously found (Pohlon et al. 2013, Gionchetta et al. 2019a, Foulquier et al. 2024, Jans et al. 2025). Previously, fungi were considered to have a greater resistance to desiccation than bacteria because their hyphal development increases their capacity to reach water or nutrients (Gionchetta et al. 2019b). Here, we found a much stronger influence of bacterial community composition than either fungal composition or diversity on decomposition, supporting recent lines of discussion that challenge the traditional vision of fungi as primary agents of decomposition (e.g., Vivanco & Martini 2025). Indeed, some bacterial groups are especially well suited to actively participate in decomposition while resisting drying. For instance, Actinobacteria are oligotrophic bacteria that grow slowly but invest heavily in extracellular enzymes, enabling them to degrade lignin, chitin, and cellulose in resource-poor environments (Delgado-Baquerizo et al. 2017, Foulquier et al. 2024). At the same time, these Gram-positive bacteria can withstand desiccation due to their thick peptidoglycan walls and ability to form extensive mycelia (Gionchetta et al. 2019a). Indeed, such bacterial groups, associated with negative values of the bacterial composite (Figure S3, Figure S8), were dominant in southern DRNs, such as GEN and BUT, where they dominated the control of decomposition rates (Figure S7). In contrast, in LEP, our northernmost DRN, bacteria and fungi also controlled decomposition rates but through completely different assemblages, dominated by clearly aquatic taxa.

These results suggest differential responses of decomposer communities to drying depending on biogeographic settings, for instance, as species from southern Europe might have developed better strategies to maintain functioning during drying since they have been exposed to this natural disturbance for much longer (Pons et al. 1995). Finally, we found strong correlations between the structure of bacterial, fungal, and even detritivore communities, a potential indicator of biotic interactions that can also shape community assembly under drying conditions (Foulquier et al. 2024). However, in our study the covariance between these three groups may also reflect commonalities in their response to drying. Future studies should investigate further the role of biotic interactions as a driver of ecosystem functioning in the context of drying river networks.

### Hydrological connectivity boosts decomposition at the network scale by reconnecting resources and consumers

At the river network scale, drying fragments the network, and thus, interrupts the flow of resources and consumers from upstream to downstream, with negative consequences for decomposition (Piano et al. 2020, Arias-Real et al. 2021). Still, high levels of connectivity after flow resumption across the network can support recovery of aquatic taxa (Sarremejane et al. 2021, Fournier et al. 2023) and functioning (Truchy et al. 2020). We found that hydrological connectivity increased decomposition rates through indirect effects associated with an enhanced contribution of fungi, bacteria, and also detritivores in non-perennial reaches. Periods of high connectivity can enhance dispersal (Jacquet et al. 2022, Chalmandrier et al. 2025) and ultimately boost decomposition through the recovery of strictly aquatic decomposers. Our analysis by DRN revealed that the main positive effects of regional connectivity on decomposition occurred in central European and northernmost networks. In contrast, connectivity had little influence on decomposition or decomposer communities in more arid, southernmost networks. These results indicate that strong levels of fragmentation can hinder their recovery of communities after flow resumption (Crabot et al. 2020, Jacquet et al. 2022), explaining long-term impacts on decomposer communities and decomposition. Furthermore, we found different responses of the various decomposer groups to connectivity, which might be associated with their different dispersal capacities and growth strategies. Fungi were the group with stronger links to regional connectivity in most SEMs. We hypothesize that increased hydrological connectivity may benefit the group with stronger dispersal limitations. Macroinvertebrate communities have mechanisms of active dispersal, which can make them less dependent on hydrological connectivity. On the other hand, the much bigger size of fungi than bacteria can limit their dispersal at regional scale (Chen et al. 2019). In contrast, bacteria, which are characterized by high diversity of physiologies and faster growth rates than fungi (Schmidt et al. 2014), are better suited to colonize and thrive in river reaches with strong environmental variations.

Besides the biotic effects, we found a positive, direct effect of hydrological connectivity on decomposition, which could be related to the redistribution across the network of important resources such as leaf litter and nutrients (Catalàn et al. 2022). In fact, the stronger direct effects of connectivity on decomposition occurred in BUK, VEL and LEP, which are characterized by intermediate levels of drying intensity but high connectivity. Thus, following Sarremejane et al. (2024), we propose that moderate levels of upstream drying may enhance downstream decomposition rates after rewetting events by delivering pulses of nutrients and high-quality leaf litter (Northington & Webster 2017, del Campo et al. 2021b, Catalàn et al. 2022). In addition, connectivity was positively correlated with discharge and water velocity in previous studies (Chalmandrier et al. 2025), which could also indicate a higher physical breakdown of leaf litter in sections with higher flow.

### Key taxa rather than diversity as drivers of decomposition in drying river networks

Drying strongly affects the diversity of macroinvertebrates (Arias-Real et al. 2021, Sarremejane et al. 2021, Chalmandrier et al. 2025, Journiac et al. 2025), bacteria and fungi (Pohlon et al. 2013, Arias-Real et al. 2023, Foulquier et al. 2024). Long dry periods can reduce decomposition through the decrease of fungal richness (Arias-Real et al. 2022). In our study, however, alpha diversity showed little effect on decomposition, except for fungi. Instead, we suggest that the main biodiversity controls of decomposition at the network scale operate through spatiotemporal changes in community composition. In previous network-scale studies, decomposition was more strongly linked to the dominance of key taxa with high decomposition ability than to diversity (Sarremejane et al. 2024, del Campo et al. 2025). Accordingly, we found the abundance of detritivores such as *Gammarus* and *Nemoura* to be important drivers of decomposition. Similarly, the composition of bacterial and fungal communities had a stronger influence on decomposition than diversity. For fungi, the effect of alpha diversity on decomposition was mainly associated with the evenness of the community rather than richness itself. Experimental studies also support a limited control of microbial diversity on decomposition under the impact of drying. Goncalves et al. (2016) reported higher decomposition at intermediate fungal diversity due to complementarity, but negative effects at high diversity from competition that inhibited efficient taxa. Furthermore, Gionchetta et al. (2024) found microbial functioning under flow intermittence to be mainly controlled by the turnover of specialist taxa, which can perform actively at different environmental conditions. Overall, our results suggest the importance of compositional changes rather than alpha diversity as the key mechanism underlying diversity-decomposition relationships in drying river networks.

### Functional implications of drying at the network scale

Recent projections of flow intermittence under climate change in European river networks found that, even under the most optimistic climate change scenarios, drying is projected to expand in space and time (Mimeau et al. 2025). Furthermore, they also found a major transition of perennial to non-perennial flow regimes, especially in Southern countries (e.g., 39 to 93 % change in BUT and GEN networks, respectively). Here, we show that short dry periods can trigger threshold responses in the diversity and composition of decomposer communities (2 to 18 dry days) and decomposition rates (50 % decline after six dry days), highlighting that just about a week of drying per year could potentially disrupt carbon cycling in river networks.

Importantly, these drying effects may interact with other global change pressures such as warming and salinization. Like drying, these two stressors can reduce the contribution of detritivores to decomposition while enhancing microbial control (Marks 2019, Canhoto et al. 2021). Interactions among these multiple stressors may have unforeseen effects for both aquatic and terrestrial food webs. For instance, while drying can exacerbate the loss of detritivores, warming may accelerate microbial decomposition, together promoting microbial dominance. Such synergistic effects among global change impacts could disrupt energy fluxes, limiting the transfer of energy into food webs through detritivores, and instead channeling it into CO₂ outgassing via microbial respiration.

Additionally, our findings position spatiotemporal drying patterns as a major regulator of biodiversity-functioning relationships at the river network scale. Besides impairing decomposition rates, the replacement of dry-sensitive taxa by generalists can also reduce functional redundancy and thus pose a risk for ecosystem resilience (Aspin et al. 2019, Leigh et al. 2019, Journiac et al. 2025). On the other hand, the loss of sensitive taxa under more severe drying scenarios might be especially detrimental for detritivore communities, where a few key taxa may control decomposition rates. Our findings underscore the role of hydrological connectivity in mitigating the impacts of drying by facilitating decomposer dispersal and restoring resource distribution. Connectivity is essential to sustain organism and resource fluxes that support natural ecosystem functioning, yet the severity of network fragmentation varies across climatic and biogeographic contexts. Moreover, how well regional connectivity may enable functional recovery after drying could also depend on spatial drying patterns (e.g. upstream vs. downstream drying; Crabot et al. 2020). Future research should further assess how the location of drying reaches within the network influences the recovery of both communities and ecosystem functioning.

## Acknowledgments

This project was financially supported by the DRYvER project, which has received funding from the European Union Horizon 2020 research and innovation program under grant agreement no. 869226. We thank Elias Dechent, Marilena Schüster, Katharina Schauer, Frédéric Boyer, Clément Lionnet, Delphine Rioux, Christian Miquel, Stephen Mulero and Dominik Gallenberger for their help with methodological aspects, field work and/or laboratory analysis.

## Notes

### Competing Interest Statement

The authors have declared no competing interest.

