## Supplemental material for "Drying halves decomposition rates in river networks by disrupting structure-function linkages"

**Section 1.** Methodological details of DNA extraction and metabarcoding procedures.

Pooled DNA purification and library preparation were performed at Fasteris facilities (Geneva, Switzerland) using the metafast PCR free protocol. A total of five libraries were prepared and then sequenced on NextSeq (2 × 150 bp, 120 m, Illumina, San Diego, USA) for Euka02 libraries, and on MiSeq (2 × 250 bp, 15 m, Illumina, San Diego, USA) for Bact02 libraries. We curated the sequencing data using the OBITools software package (Boyer et al. 2016) and self-made R scripts (Taberlet et al. 2018). In brief, the paired-end reads were assembled and assigned to their respective samples/markers before being dereplicated. Low-quality sequences were excluded. Pairwise dissimilarities were computed between sequences using the Sumatra algorithm (Mercier et al. 2013) with a maximum of 3 mismatches allowed. MOTUs were then formed by clustering sequences at 97% similarity using the Sumaclust algorithm (Mercier et al. 2013). The abundance of a MOTU was defined as the sum of the read abundances of the sequences belonging to it. In subsequent analyses, each MOTU was represented by its most abundant sequence. Each MOTU was assigned to a taxonomic clade with the ecotag program (Boyer et al. 2016). We retained only taxonomic annotations with >80% identities. We removed MOTUs peaking in abundance in blank extraction or PCR negative controls. Furthermore, we removed PCR replicates with a number of reads and MOTU lower or similar to PCR negative controls (Taberlet et al. 2018). At the end of this process, PCR replicates were pooled for each site.

**Section 2.** R code used to build the four structural equation models tested in the study based on the Lavaan package. STcon = Spatiotemporal connectivity; bac.comp = bacterial composite; fun.comp = fungal composite; det.abund = Detritivore abundance; det.comp = detritivore composite. DRN = drying river network.

```
# SEM1: Full SEM with detritivore abundance

SEM1 <- '
#Regressions
k ~ drydays + STcon + bac.comp + fun.comp + det.abund
bac.comp ~ drydays + STcon
fun.comp ~ drydays + STcon
det.abund ~ drydays + STcon
# correlation errors
bac.comp ~~ fun.comp
bac.comp ~~ det.abund
'
SEM1.fit = sem(SEM1, data=df, fixed.x = FALSE)
summary(SEM1.fit, standardize = T, rsq=T, fit.measures = T)
modindices(SEM1.fit, sort=T)

# SEM1.1: SEM1 with only flowing reaches and the detritivore composite

SEM1.1 <- '
#Regressions
k ~ drydays + STcon + bac.comp + fun.comp + det.comp
bac.comp ~ drydays + STcon
fun.comp ~ drydays + STcon
det.comp ~ drydays + STcon
# correlation errors
bac.comp ~~ fun.comp
bac.comp ~~ det.comp
'
SEM1.1.fit = sem(SEM1.1, data=df, fixed.x = FALSE)
summary(SEM1.1.fit, standardize = T, rsq=T, fit.measures = T)
modindices(SEM1.1.fit, sort=T)

# SEM2: Multigroup SEM with flow regime as grouping variable and detritivore abundance

SEM2 <- '
#Regressions
# Non-perennial
group: 1
k ~ drydays + STcond30 + bac.comp + fun.comp + shrF.abundC
bac.comp ~ drydays + STcond30
fun.comp ~ drydays + STcond30
shrF.abundC ~ drydays + STcond30
# correlation error between covariates
```

```
bac.comp ~~ fun.comp
bac.comp ~~ shrF.abundC
# Perennial
group: 2
k ~ STcond30 + bac.comp + fun.comp + shrF.abundC
bac.comp ~ STcond30
fun.comp ~ STcond30
shrF.abundC ~ STcond30
# correlation error between covariates
bac.comp ~~ fun.comp
bac.comp ~~ shrF.abundC
'
SEM2.fit = sem(SEM2, data = df, group = "Regime", fixed.x = F)
summary(SEM2.fit, standardize = T, rsq=T, fit.measures = T)
modindices(SEM2.fit, sort=T)


# SEM3: Multigroup SEM with river network identity as grouping variable and detritivore abundance

SEM3 <- '
#Regressions
k ~ drydays + STcon + bac.comp + fun.comp + det.abund
bac.comp ~ drydays + STcon
fun.comp ~ drydays + STcon
det.abund ~ drydays + STcon
# correlation error
bac.comp ~~ fun.comp
bac.comp ~~ det.abund
'
SEM3.fit = sem(SEM3, data = df, group = "DRN", fixed.x = F)
summary(SEM3.fit, standardize = T, rsq=T, fit.measures = T)
modindices(SEM3.fit, sort=T)
```

**Table S1.** Summary of the temporal overlap between the leaf litter decomposition experiments (DE) and biodiversity sampling campaigns. Starting (date0) and final dates (dateF) for both campaigns are provided as well as the number of flowing sites (F), dry sites (D / DP) and the percentage of flowing sites (%F) during each campaign. DRN = drying river network.

| | | WP3 / Decomposition experiment | | | | | WP2 / Biodiversity sampling | | | | | |
|---|---|---|---|---|---|---|---|---|---|---|---|---|
| DRN* | Camp | date0 | dateF | F | D/DP | %F | Camp | date0 | dateF | F | D/DP | Temporal overlap |
| ALB | 1 | 16/03/21 | 22/04/21 | 20 | 0 | 100 | 1 | 16/03/21 | 01/04/21 | 17 | 0 | during DE |
| ALB | 2 | 22/06/21 | 21/07/21 | 17 | 2 | 89 | 3 | 21/06/21 | 02/07/21 | 17 | 2 | during DE |
| ALB | 3 | 25/10/21 | 03/12/21 | 17 | 3 | 85 | 5 | 22/11/21 | 14/12/21 | 10 | 2 | during DE |
| BUK | 1 | 16/03/21 | 17/04/21 | 19 | 1 | 95 | 2 | 19/04/21 | 29/04/21 | 20 | 0 | 2-12 days after DE |
| BUK | 2 | 14/07/21 | 29/07/21 | 12 | 8 | 60 | 4 | 10/08/21 | 17/08/21 | 9 | 9 | 12-19 days after DE |
| BUK | 3 | 12/10/21 | 01/12/21 | 15 | 5 | 75 | 6 | 13/12/21 | 21/12/21 | 17 | 5 | 12-20 days after DE |
| BUT | 1 | 15/03/21 | 27/04/21 | 16 | 3 | 84 | 2 | 20/04/21 | 04/05/21 | 13 | 5 | during DE |
| BUT | 2 | 22/06/21 | 21/07/21 | 12 | 9 | 57 | 4 | 13/07/21 | 14/07/21 | 10 | 9 | during DE |
| BUT | 3 | 22/09/21 | 20/11/21 | 16 | 4 | 80 | 6 | 10/11/21 | 16/11/21 | 12 | 9 | during DE |
| GEN | 1 | 27/03/21 | 29/05/21 | 20 | 0 | 100 | 2 | 21/03/21 | 26/03/21 | 20 | 0 | 1-6 days before DE |
| GEN | 2 | 13/07/21 | 27/07/21 | 12 | 8 | 60 | 4 | 23/07/21 | 27/07/21 | 10 | 8 | during DE |
| GEN | 3 | 26/10/21 | 22/01/22 | 17 | 3 | 85 | 6 | 06/02/22 | 12/02/22 | 16 | 3 | 15-21 days after DE |
| LEP | 1 | 17/05/21 | 11/06/21 | 20 | 0 | 100 | 3 | 14/06/21 | 17/06/21 | 20 | 0 | 3-6 days after DE |
| LEP | 2 | 18/08/21 | 16/09/21 | 15 | 5 | 75 | 4 | 09/08/21 | 12/08/21 | 15 | 6 | 6-9 days before DE |
| LEP | 3 | 07/10/21 | 07/11/21 | 20 | 0 | 100 | 6 | 01/11/21 | 03/11/21 | 20 | 0 | during DE |
| VEL | 1 | 04/05/21 | 27/05/21 | 20 | 0 | 100 | 2 | 04/05/21 | 05/05/21 | 19 | 0 | during DE |
| VEL | 2 | 19/07/21 | 07/08/21 | 12 | 8 | 60 | 4 | 17/08/21 | 22/08/21 | 12 | 7 | 10-15 days after DE |
| VEL | 3 | 11/10/21 | 15/11/21 | 13 | 7 | 65 | 6 | 23/11/21 | 24/11/21 | 12 | 10 | 8-9 days after DE |

* GEN = Genal (Spain), BUT = Butižnica (Croatia), ALB = Albarine (France), BUK = Bükkösdi-víz (Hungary), VEL = Velička (Czech Republic), LEP = Lepsämänjoki (Finnland)

.

**Table S2.** Summary of the drying patterns across the six studied drying river networks (DRN). Dry days were estimated during the periods of the decomposition experiments (Table S1), while the hydrological connectivity was computed for a temporal window of 30 days before the start of the experiments. Hydrological connectivity varies from 0 to 1, indicating changes in connectivity from total disconnection across the network to fully connected. Values of dry days correspond only to non-perennial reaches (NP) at each network, while values for hydrological connectivity are computed for all reaches. DRN = drying river network.

| | Number of dry days (NP) | | Hydrological connectivity | |
|---|---|---|---|---|
| DRN* | median | range | median all | range |
| GEN | 7 | 0-50 | 0.07 | 0-0.78 |
| BUT | 43 | 0-66 | 0.13 | 0-1 |
| ALB | 5 | 0-24 | 0.45 | 0.20-0.97 |
| BUK | 11 | 0-51 | 0.23 | 0-1 |
| VEL | 15 | 0-35 | 0.24 | 0-1 |
| LEP | 0 | 0-4 | 0.98 | 0-1 |

* GEN = Genal (Spain), BUT = Butižnica (Croatia), ALB = Albarine (France), BUK = Bükkösdi-víz (Hungary), VEL = Velička (Czech Republic), LEP = Lepsämänjoki (Finnland).

**Table S3.** Top-performing linear models (delta BIC < 2) identifying the best combination of predictors for explaining decomposition rates based on the diversity and composition of each decomposer group. Models marked in bold were chosen as the best-performing models. Model selection was based on BIC (Bayesian Information Criterion) to promote more parsimonious models. df = degrees of freedom; RMSE = root mean squared error; AIC = Akaike Information Criterion; abund = abundance; rich = richness; Piel = Pielou evenness.

| | Standardized coefficients | | | | | Model performance | | | | | |
|---|---|---|---|---|---|---|---|---|---|---|---|
| Bacteria | | | | | | | | | | | |
| | bac. pcoa1 | bac. pcoa2 | bac. pcoa3 | bac. pcoa6 | bac. pcoa8 | df | RMSE | AIC | BIC | ΔBIC | R2 |
| **1** | **0.266** | **0.185** | **-0.322** | **-0.112** | **0.093** | **7** | **0.418** | **312** | **336.8** | **0** | **0.51** |
| Fungi | | | | | | | | | | | |
| | fun. pcoa1 | fun. pcoa2 | fun. pcoa5 | Piel. | rich. | df | RMSE | AIC | BIC | ΔBIC | R2 |
| **1** | **-0.192** | **0.247** | | **0.215** | **-0.121** | **6** | **0.5** | **395.6** | **417** | **0** | **0.30** |
| 2 | -0.188 | 0.250 | 0.066 | 0.217 | -0.148 | 7 | 0.497 | 393.6 | 418.6 | 1.55 | 0.31 |
| Detritivores* | | | | | | | | | | | |
| | abund. | det. pcoa5 | det. pcoa6 | det. pcoa7 | rich. | df | RMSE | AIC | BIC | ΔBIC | R2 |
| 1 | 0.144 | -0.080 | -0.071 | 0.092 | | 6 | 0.439 | 282.6 | 303.1 | 0 | 0.18 |
| **2** | **0.159** | **-0.080** | | **0.087** | | **5** | **0.444** | **286.1** | **303.2** | **0.06** | **0.17** |
| 3 | 0.118 | | | 0.119 | 0.086 | 5 | 0.446 | 287.5 | 304.6 | 1.52 | 0.16 |
| 4 | 0.159 | | | 0.088 | | 4 | 0.452 | 291.2 | 304.9 | 1.83 | 0.15 |
| 5 | 0.144 | | -0.071 | 0.093 | | 5 | 0.446 | 287.9 | 305 | 1.9 | 0.16 |
| * Model selection for detritivores only included observations from flowing reaches. | | | | | | | | | | | |

**Table S4.** Comparison of model performance for the four structural equation models (SEM). The main indicator of goodness of fit for SEM using “lavaan” are chi2 and p-values. Lower chi2 values indicate better performance, whereas p values > 0.05 indicate that the proposed model has no discrepancies with the observed data. Other measurements of model fit that indicate good performance are the root-mean squared error of approximation (RMSEA) < 0.08, the comparative fit index (CFI) > 0.9, or the standardized root-mean squared residual (SRMR) <0.08. n = sample size; NP = non-perennial reaches, P = perennial reaches. DRN = drying river network.

| | SEM1 | SEM1.1 | SEM2 | SEM3 |
|---|---|---|---|---|
| Description | Flowing & dry reaches, detritivore abundance (Fig. 5) | Modification SEM1: only flowing reaches, detritivore composite (Fig. S6) | Multigroup SEM with flow regime as grouping factor (Fig. 6) | Multigroup SEM with DRN as grouping factor (Fig. S7) |
| n | 262 | 214 | NP: 127, P: 135 | GEN: 42, BUT: 28, ALB: 43, BUK: 51, VEL:49, LEP:49 |
| chi2 | 1.591 | 0.048 | 4.494 (NP: 4.341, P:0.153) | 12.006 GEN: 1.373, BUT: 0.001, ALB: 7.422 BUK: 0.013., VEL:1.688, LEP:1.508 |
| df | 1 | 1 | 2 | 6 |
| p value | 0.207 | 0.826 | 0.106 | 0.062 |
| k $R^2$ | 0.62 | 0.40 | NP: 0.67, P: 0.29 | GEN: 0.78, BUT: 0.81, ALB: 0.32, BUK: 0.65, VEL:0.68, LEP: 0.37 |
| AIC | 3131 | 1964 | 2711 | 2645 |
| BIC | 3203 | 2031 | 2872 | 3201 |
| CFI | 0.999 | 1 | 0.995 | 0.992 |
| RMSEA | 0.047 | 0 | 0.098 | 0.151 |
| SRMR | 0.011 | 0.003 | 0.015 | 0.020 |

* GEN = Genal (Spain), BUT = Butižnica (Croatia), ALB = Albarine (France), BUK = Bükkösdi-víz (Hungary), VEL = Velička (Czech Republic), LEP = Lepsämänjoki (Finnland)

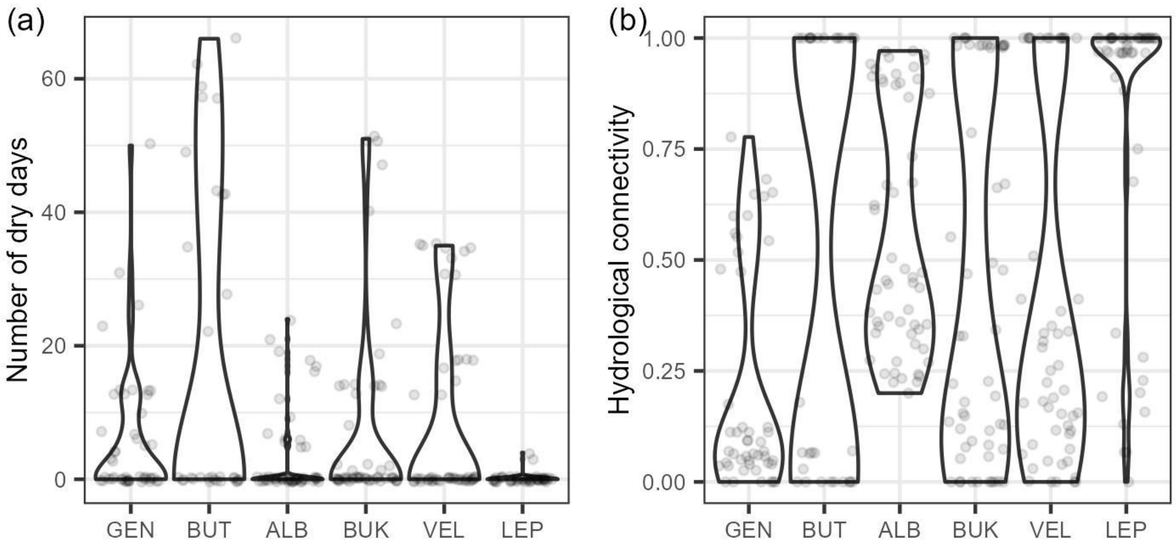


**Figure S1.** Violin plots representing differences in (a) number of dry days and (b) hydrological connectivity across drying river networks. Networks are distributed along the x axis following a latitudinal gradient from south (left) to north (right). GEN = Genal (Spain), BUT = Butižnica (Croatia), ALB = Albarine (France), BUK = Bükkösdi-víz (Hungary) , VEL = Velička (Czech Republic), LEP = Lepsämänjoki (Finnland).

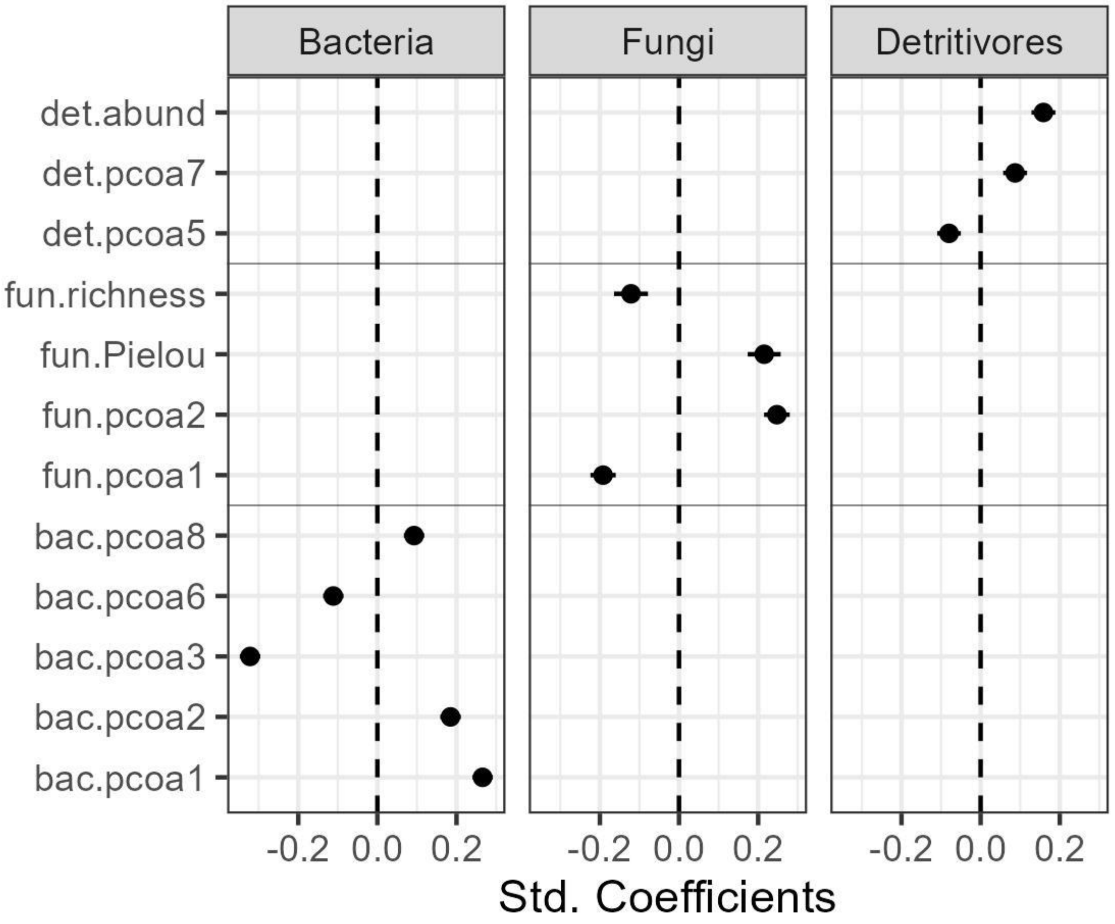


**Figure S2.** Standardized (Std.) coefficients (mean ± 95% confidence interval) of predictors extracted from the best linear models explaining decomposition rates obtained independently for each decomposer group. Selected predictors were used to create composite variables used in SEMs to characterize the role of decomposer communities in litter decomposition. See interpretation of predictors in Table 1.

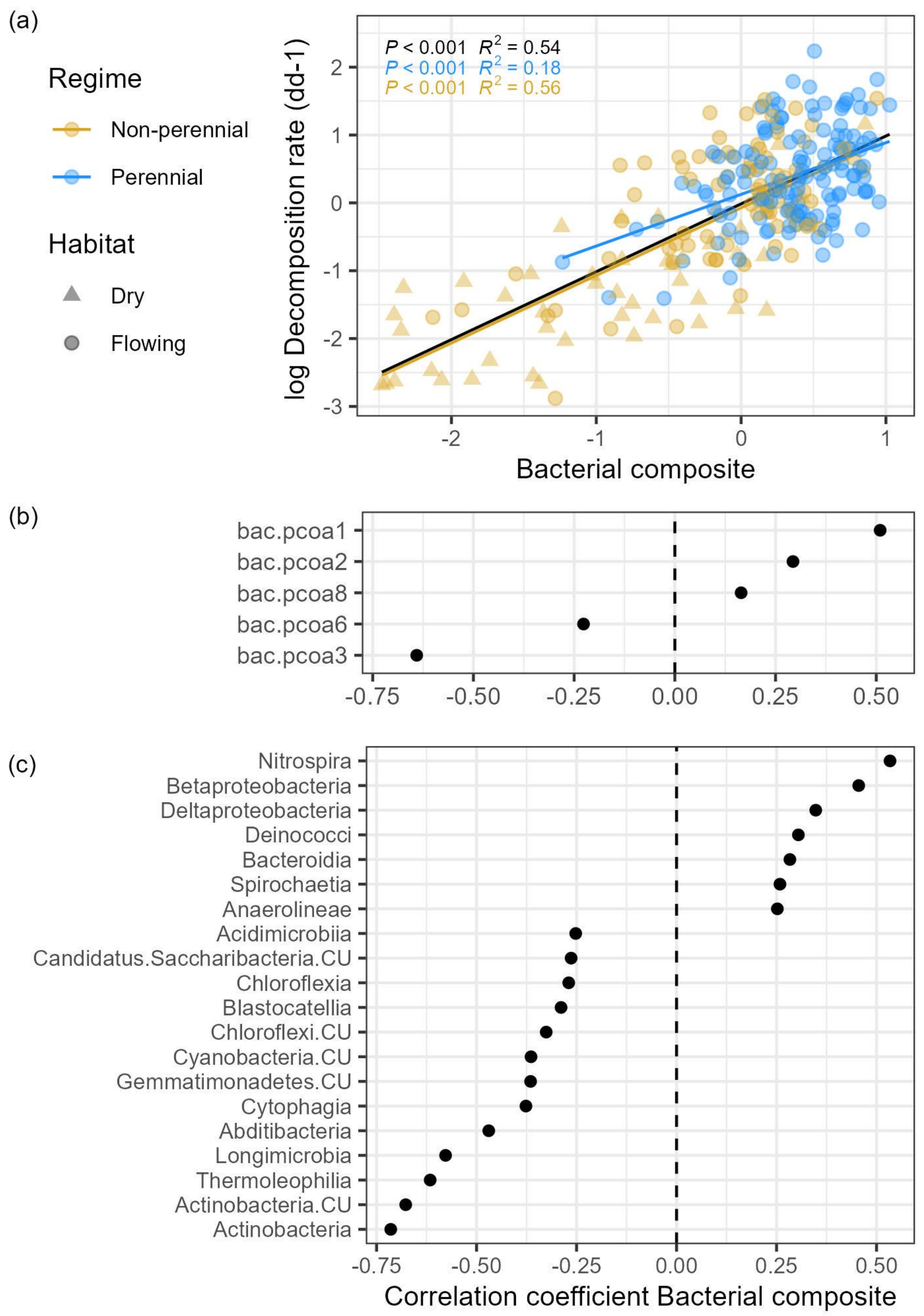


**Figure S3.** Construction of bacterial composite used in SEM to summarise the effect of bacterial community composition on decomposition (a). The composite arises from a multiple regression model that considers the combined effect of various PCoA components describing bacterial community composition on decomposition rates (Table S3). The black line represents the linear fit between decomposition rates and the bacterial composite when using all observations; blue and orange lines represent the linear fit for each flow regime. Correlation plots (b and c) show the direction and magnitude of the correlation between the bacterial composite and (b) the selected predictors, or (c) related bacteria MOTUS grouped by class. CU = class undefined.

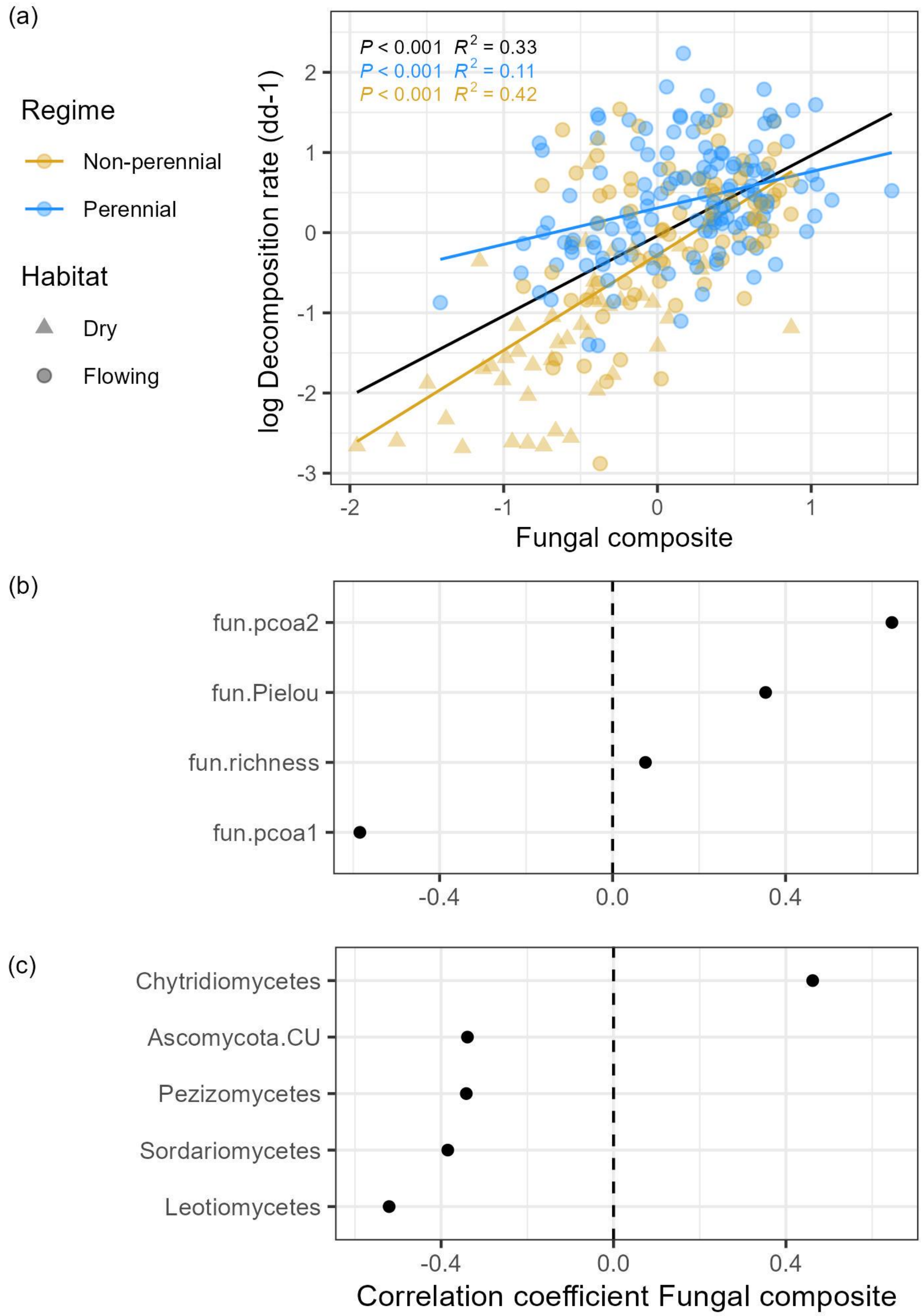


**Figure S4.** Construction of fungal composite used in SEM to summarise the effect of fungal community composition and diversity on decomposition (a). The composite arises from a multiple regression model that considers the combined effect of fungal richness, evenness and various PCoA components describing fungal community composition on decomposition rates (Table S3). The black line represents the linear fit between decomposition rates and the fungal composite when using all observations; blue and orange lines represent the linear fit for each flow regime. Correlation plots (b and c) show the direction and magnitude of the correlation between the fungal composite and (b) the selected predictors, or (c) related fungi MOTUS grouped by class. CU = class undefined.

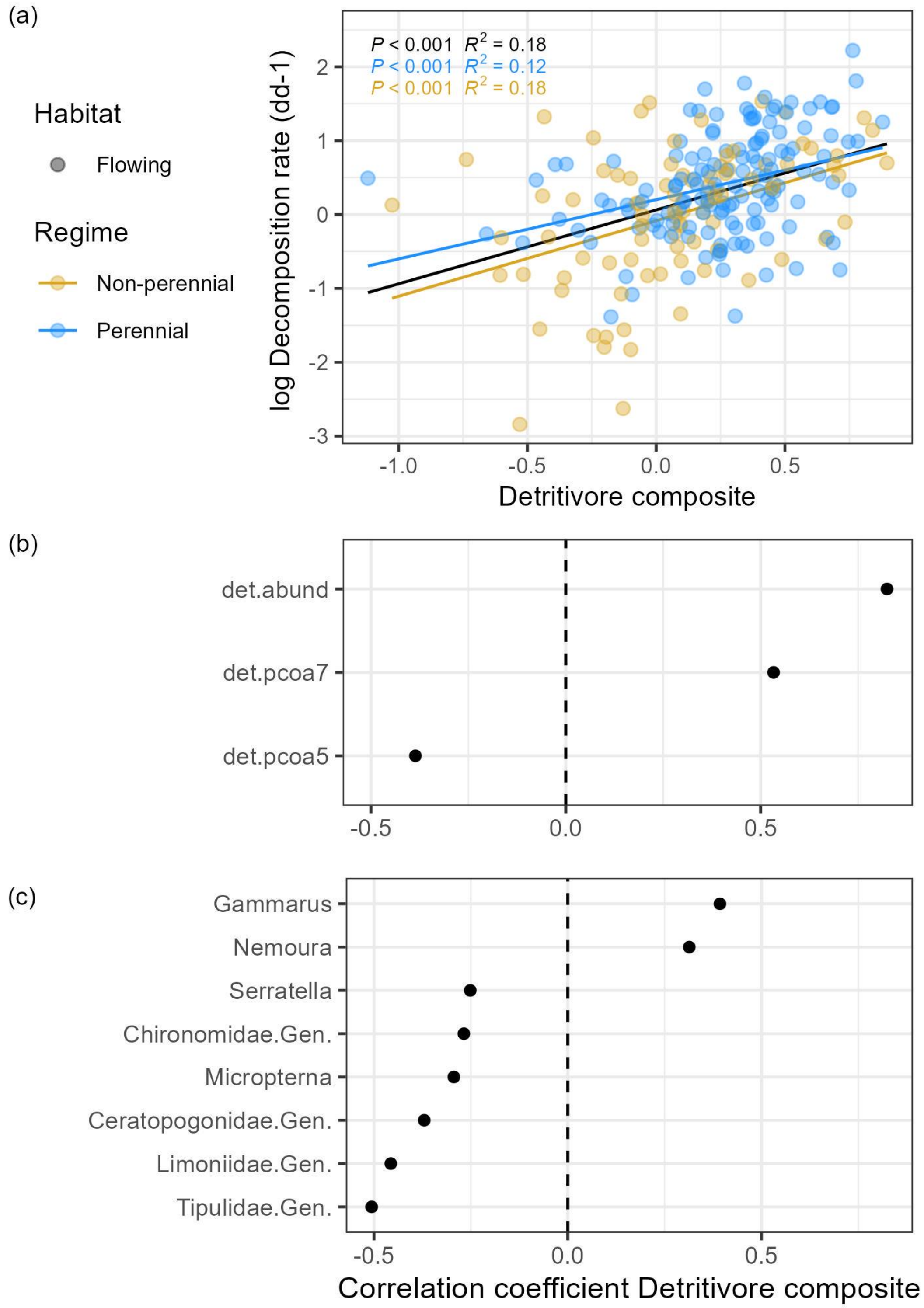


**Figure S5.** Construction of the detritivore composite used in SEM1.1 to summarise the effect of detritivore abundance and community composition on decomposition (a). The composite arises from a multiple regression model that considers the combined effect of total detritivore abundance and various PCoA components describing detritivore community composition on decomposition rates (Table S3). The black line represents the linear fit between decomposition rates and the detritivore composite when using all observations; blue and orange lines represent the linear fit for each flow regime. Correlation plots (b and c) show the direction and magnitude of the correlation between the detritivore composite and (b) the selected predictors, or (c) related detritivore taxa.

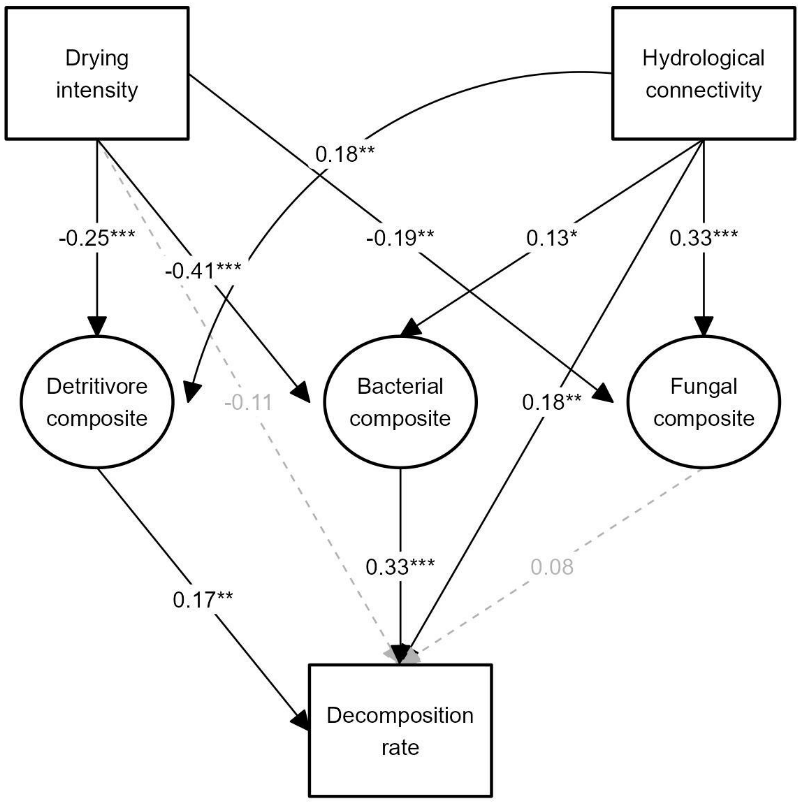


**Figure S6.** Alternative version of SEM1 (SEM1.1) to evaluate long-term effects of drying on decomposition in flowing conditions through changes in detritivore community composition (summarised by the detritivore composite, Figure S5). Boxes represent measured parameters, while the circles represent composite variables. Black solid lines and associated numbers represent significant paths and standardised path coefficients. Grey discontinuous lines and associated numbers represent non-significant paths.

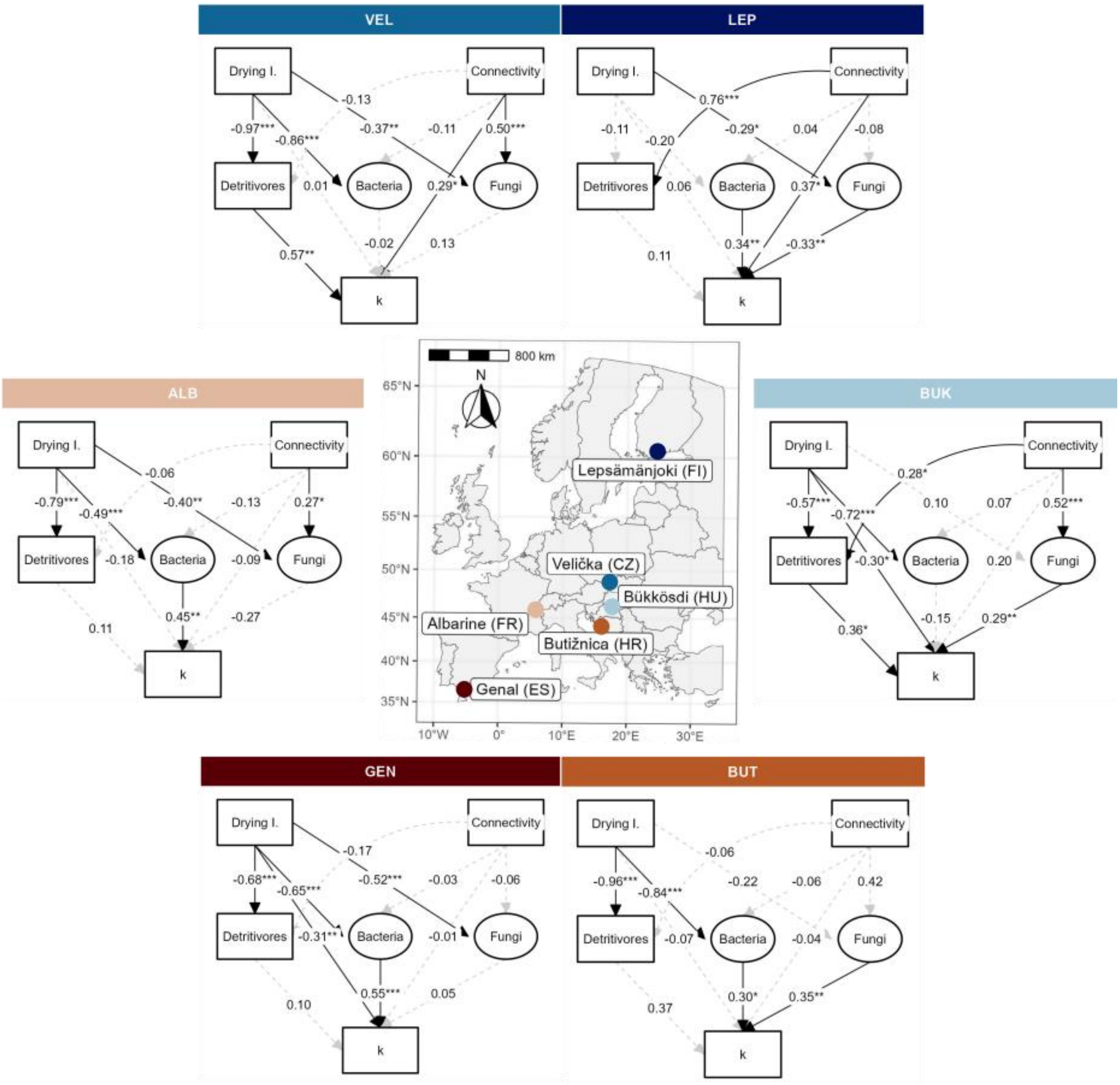


**Figure S7.** Multigroup SEM describing differential biotic controls of decomposition rates depending on the different drying river networks (SEM3). SEM graphs are distributed following a latitudinal gradient from south (bottom) to north (top). Boxes represent measured parameters, while the circles represent composite variables created to summarise the effect of bacterial and fungal community composition on litter decomposition. Solid lines and associated numbers represent significant paths and standardised path coefficients. Discontinuous lines and associated grey numbers represent non-significant paths. Drying I. = drying intensity, Connectivity = hydrological connectivity, Detritivores = Detritivore abundance, Bacteria = bacterial composite, Fungi = Fungal composite, k = decomposition rates.

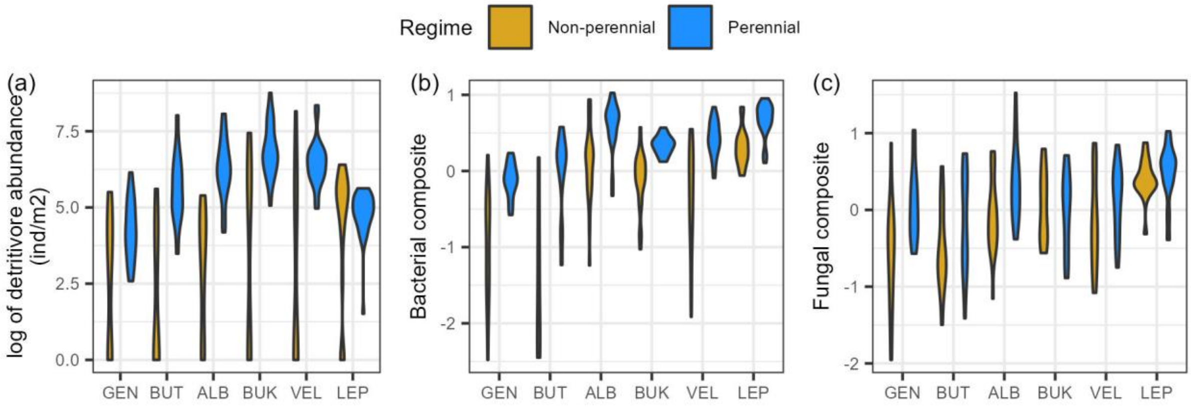


**Figure S8.** Violin plots representing differences in (a) the abundance of detritivores, (b) the bacterial composite, and (c) the fungal composite across drying river networks and among flow regimes. Networks are distributed along the x axis following a latitudinal gradient from south (left) to north (right). GEN = Genal (Spain), BUT = Butižnica (Croatia), ALB = Albarine (France), BUK = Bükkösdi-víz (Hungary), VEL = Velička (Czech Republic), LEP = Lepsämänjoki (Finnland).
